# The gases H_2_ and O_2_ in open-culture reactors influence the performance and microbiota of chain elongation into *n*-caproate and *n*-caprylate

**DOI:** 10.1101/2024.03.21.586105

**Authors:** Catherine M. Spirito, Timo N. Lucas, Sascha Patz, Byoung Seung Jeon, Jeffrey J. Werner, Lauren H. Trondsen, Juan J. Guzman, Daniel H. Huson, Largus T. Angenent

## Abstract

Medium-chain carboxylates are used in various industrial applications. These chemicals are typically extracted from palm oil, which is deemed not sustainable. Recent research has focused on microbial chain elongation using reactors to produce medium-chain carboxylates, such as *n*-caproate (C6) and *n*-caprylate (C8), from organic substrates such as wastes. Even though the production of *n*-caproate is relatively well-characterized, bacteria and metabolic pathways that are responsible for *n*-caprylate production are not. Here, three 5-L reactors with continuous membrane-based liquid-liquid extraction (*i.e.*, pertraction) were fed ethanol and acetate and operated for an operating period of 234 days with different operating conditions. Metagenomic and metaproteomic analyses were employed. *n*-Caprylate production rates and reactor microbiomes differed between reactors even when operated similarly due to differences in H_2_ and O_2_ between the reactors. The complete reverse β-oxidation pathway was present and expressed by several bacterial species in the *Clostridia* class. Several *Oscillibacter* spp., including *Oscillibacter valericigenes*, were positively correlated with *n*-caprylate production rates, while *Clostridium kluyveri* was positively correlated with *n*-caproate production. *Pseudoclavibacter caeni*, which is a strictly aerobic bacterium, was abundant across all the operating periods, regardless of *n*-caprylate production rates. This study provides insight into microbiota that are associated with *n*-caprylate production in open-culture reactors and provides ideas for further work.

**Importance:** Microbial chain elongation pathways in open-culture biotechnology systems can be utilized to convert organic waste and industrial side streams into valuable industrial chemicals. Here, we investigated the microbiota and metabolic pathways that produce medium-chain carboxylates, including *n*-caproate (C6) and *n*-caprylate (C8), in reactors with in-line product extraction. Although the reactors in this study were operated similarly, different microbial communities dominated and were responsible for chain elongation. We found that different microbiota were responsible for *n*-caproate or *n*-caprylate production, and this can inform engineers on how to operate the systems better. We also observed which changes in operating conditions steered the production toward and away from *n*-caprylate, but more work is necessary to ascertain a mechanistic understanding that could be predictive. This study provides pertinent research questions for future work.

## Introduction

Medium-chain carboxylates (MCCs), such as *n*-caproate and *n*-caprylate, are utilized in a variety of industrial and agricultural applications, including as biofuel precursors, anticorrosion agents, plasticizers, personal care products, feed additives, and antimicrobials (1). MCCs are typically produced as a byproduct of palm-oil refining (2). Recent research has focused on producing MCCs in open-culture reactors from organic substrates, including wastes, as part of a circular economy. MCCs have a relatively low solubility in water in their undissociated form. Therefore, MCCs can be extracted from aqueous broths *via* techniques, such as in-line product extraction, to address MCC toxicity issues and to increase volumetric production rates (3, 4). Laboratory studies demonstrated the efficient production of MCCs by anaerobic fermenter microbiomes at rates comparable to methane production by anaerobic digester microbiomes (4, 5). MCCs are produced *via* pure and open cultures from a variety of substrates, including: **(1)** synthetic substrates utilizing ethanol (3, 6, 7) or lactic acid (8, 9) as the electron donor; **(2)** organic wastes; and **(3)** industrial side streams (10-18).

Medium-chain carboxylates are often produced *via* the reverse β-oxidation (RBOX) pathway in which ethanol, lactic acid, or another electron donor is oxidized to acetyl-CoA *via* substrate-level phosphorylation by alcohol dehydrogenase and acetaldehyde dehydrogenase. This provides the energy (ATP) and reducing equivalents for chain elongation of short-chain carboxylates, such as acetate and *n*-butyrate, to longer-chain carboxylates, such as *n*-caproate (six carbon chain) and *n*-caprylate (eight carbon chain) (1, 19-21) (**Fig. 1**). Chain elongation is a cyclic process in which acetyl-CoA enters the cycle and is condensed with an acyl-CoA by acetyl-CoA C-acyltransferase (Thiolase II) (ACAT) to form an acyl-CoA that is two carbon atoms longer than its substrate. The product of this reaction is further reduced by 3-hydroxy-acyl-CoA dehydrogenase (HAD) or 3-hydroxy-butyrl-CoA dehydrogenase (HBD) (**Fig. 1**). This product is then dehydrated to 2-enoyl-CoA by enoyl-CoA dehydratase (ECH) and further reduced by an acyl-CoA dehydrogenase (ACD) or butryl-CoA dehydrogenase to form an elongated acyl-CoA (**Fig. 1**). Finally, terminal enzymes acetyl-CoA transferase (CoAT) or thioesterase (TE) act to remove coenzyme A from the terminal acyl-CoA molecule and release the corresponding acid (**Fig. 1**). Energy is conserved during the RBOX pathway *via* flavin-based electron bifurcation and the Rnf respiratory complex (RNF) in which an electron-bifurcating acyl-CoA dehydrogenase complex utilizes two electron-transfer flavoproteins (**Fig. 1**) (22). *Prior* research primarily focused on RBOX as the pathway for MCC production (4, 7, 23-25). A few recent studies have suggested that the fatty acid biosynthesis (FAB) pathway may play a role as well (26, 27), though, it should be noted that this pathway is used by all bacteria to build their phospholipid membranes. In addition, FAB is an anabolic process, it consumes energy rather than producing energy that is necessary for bacterial growth in anaerobic conditions without inorganic electron acceptors for respiration.

**Figure 1:**
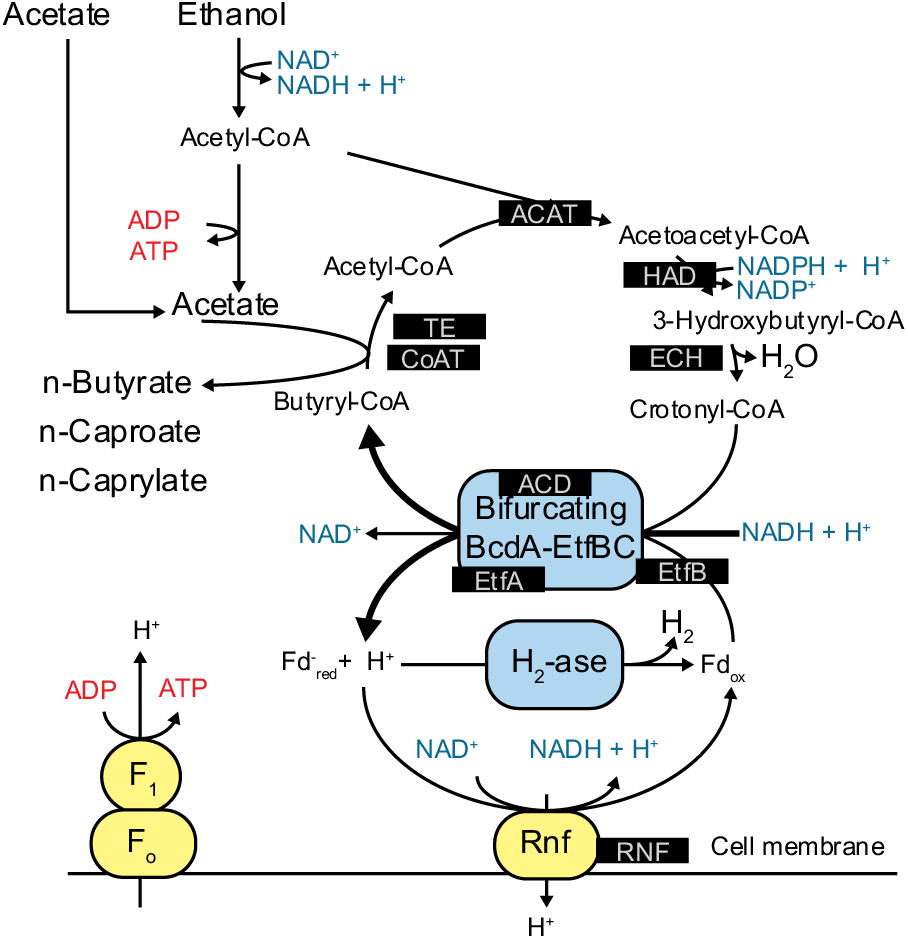
The reverse β-oxidation (RBOX) pathway that was investigated in this study. The enzymes we examined in this study are highlighted in black boxes. The figure was modified with permission from Angenent et al. 2016 (21). RBOX pathway enzymes are: ACAT: acetyl-CoA C-acyltransferase (Thiolase II); HAD: 3-hydroxy-acyl-CoA dehydrogenase; ECH: enoyl-CoA dehydratase; ACD: acyl-CoA dehydrogenase EtfA/B: electron-transfer-flavoprotein subunit A/B; CoAT: Acetyl CoA-transferase; TE: Thioesterase; RNF: Rnf respiratory complex.

Previously, both pure- and open-culture studies identified multiple bacterial strains that produce *n*-caproate. Known *n*-caproate-producing bacteria primarily belong to the phylum Firmicutes, except for *Rhodospirillum rubrum* (28). Within the phylum Firmicutes, *n*-caproate-producing bacterial strains have been isolated and identified from six genera: *Caproicibacter*, *Caproiciproducens*, *Clostridium* (29-31), *Eubacterium* (32, 33), *Megasphaera* (34), and *Pseudoramibacter* (24). In open-culture reactor studies, certain bacteria have been associated with high *n*-caprylate production rates, including *Burkholderia* spp., *Clostridium* group IV spp., *Desulfosporosinus meridiei*, *Oscillospira* spp., Rhodocyclaceae K82 spp., unknown Ruminococcaceae, and *Sphingobacterium multiform* (3, 4, 11). These studies were based on 16S rRNA gene sequencing data. Few bacterial isolates have been shown to produce *n*-caprylate. This is attributed to the microbial toxicity of *n*-caprylate, and the lack of measurement of *n*-caprylate in *prior* studies. In 1967, *Ramibacterium alactolyticum*, which has been renamed to *Pseudoramibacter alactolyticus*, was shown to produce *n*-caproate and *n*-caprylate from glucose (35). A previous pure-culture reactor study observed the production of relatively low concentrations of *n*-caprylate by *Clostridium kluyveri*, which is a well-known chain-elongating bacterium, in a reactor fed a 10:1 molar ratio of ethanol and acetate (*i.e.*, syngas effluent), operated at pH 7 and with an in-line membrane-based liquid/liquid extraction (*i.e.,* pertraction) system to reduce the toxicity (10). At lower pH levels, a lower rate of *n*-caprylate production was observed. For the open-culture studies, a shotgun metagenomics study found an uncultured Clostridiales order bacterium, *Candidatus Weimeria bifida*, gen. nov., sp. nov., which could produce *n*-caprylate from xylose (23, 24). Research is needed to understand essential players and metabolic pathways in these reactors that optimize *n*-caprylate production.

Here, we investigated the role of the RBOX pathway in producing *n*-caproate and *n*-caprylate. Our original objective was to build and operate three equal stain-less steel reactor systems to prevent O_2_ intrusion as much as possible. As an independent operating unit, we planned different H_2_ concentrations for each of the three systems. However, we show here that we were not successful in: **(1)** preventing O_2_ intrusion; and **(2)** utilizing H_2_ as an independent parameter. Regardless, we obtained pertinent data by operating the three 5-L open-culture reactors with in-line product extraction throughout 234 days. We employed Illumina 16S rRNA gene sequencing, shotgun metagenomics, and metaproteomics to characterize microbiomes. The reactors were fed ethanol and acetate and produced *n*-caproate and *n*-caprylate. Even though the reactors were all provided the same substrates and produced MCCs, their microbiota differed. Several bacterial species belonging to the class Clostridia, including *Oscillibacter valericigenes*, expressed the majority of the RBOX pathway. *Oscillibacter* spp. members were found to be positively correlated with *n*-caprylate production rates. The aerobic bacterium *Pseudoclavibacter caeni* was one of the abundant bacteria in the reactor samples, regardless of *n*-caprylate production rates. *P. caeni* may have acted as an O_2_ scavenger in the system or provided other unknown roles for producing *n*-caprylate.

## Results

### The performance of the three reactors differed despite similar operating conditions

We operated three stainless-steel, continuously stirred reactors with a 5-L working volume and in-line product extraction at mesophilic conditions and a pH of 5.5 (**Table 1, Fig. S1**). After a 75-day start-up period and at the start of Period I, we mixed the microbiota from all three reactors and then operated the three reactors similarly throughout Period I (without sparging). Regardless, the performance of the three reactors was not similar and varied during this period. Reactors 1 and 2 achieved promising and similar overall medium-chain production rates (**Fig. 2**), but Reactor 3 performed poorly during Period 1 with a lower *n*-caprylate production rate compared to Reactors 1 and 2 (**Fig. 2**). Reactor 1 exhibited a higher selectivity (*i.e.*, wanted products compared to the substrate) for *n*-caprylate production compared to Reactor 2. The maximum average volumetric *n*-caprylate production rate was 1.1×10^2^ ± 7.1 mmol C L^-1^ d^-1^ (0.080 ± 0.005 g L^-1^ h^-1^) during Period 1D for Reactor 1 (**Fig. 2B**). Small differences in operating conditions, potentially due to differences in the tightness of the reactor seals resulting in different H_2_ and O_2_ exchange conditions, seem to have had an amplified impact on chain elongation. This was different from anaerobic digestion of animal waste for which we found that four similar reactor operating conditions resulted in almost identical performances after an operating period of one year (36).

**Figure 2.**
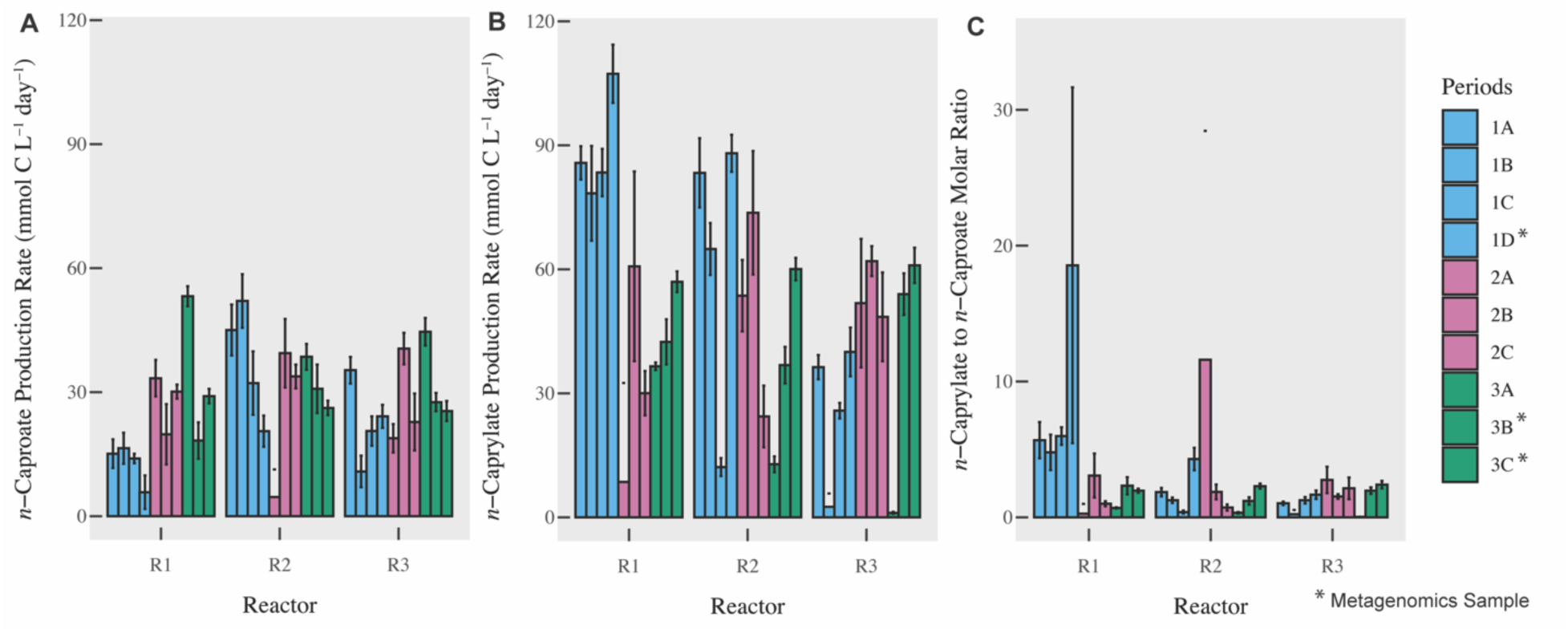
*n*-caproate (**A**) and *n*-caprylate (**B**) total production rates (mmol C L^-1^ d^-1^) and the molar ratio of *n*-caprylate to *n*-caproate (**C**) in three reactors across three main periods (and ten periods). Period divisions are explained in the methods. Error bars indicate the standard error for the measurements. *Legend indicates periods (Periods 1D, 3B, 3C) in which biomass samples were collected from reactors for shotgun metagenomic analysis. R1-3 are Reactors 1-3.

**Table 1.** Operating data for three reactors. We report the hydraulic retention time (HRT), organic loading rate (OLR), and gas flow rate for each reactor and each period. The gas flow rate was measured at the outlet of each reactor system. Different gases were utilized as indicated. During Period 2, gas sparging was periodically on and off to the reactors, whereas it was on all the time during Period 3. Mean ± s.e. is reported. R1-3 are Reactors 1-3.

| Reactor | Period | Days | HRT (d) | OLR<br>(mmol C L<sup>-1</sup> d<sup>-1</sup>) | Gas Flow Rate<br>(L d<sup>-1</sup>) | Sparged with |
| --- | --- | --- | --- | --- | --- | --- |
| R1 | 1 | 75 to 142 | 8.9 ± 0.3 | 1.4x10<sup>2</sup> ± 7.9 | 0 | No gas |
| | 2 | 143 to 184 | 9.6 ± 0.3 | 1.2x10<sup>2</sup> ± 6.7 | 3.38±4.86 | N<sub>2</sub> off/on |
| | 3 | 185 to 234 | 9.0 ± 0.3 | 1.4x10<sup>2</sup> ± 6.5 | 24.7±13.7 | N<sub>2</sub> |
| R2 | 1 | 75 to 142 | 8.9 ± 0.5 | 1.4x10<sup>2</sup> ± 8.4 | 0 | No gas |
| | 2 | 143 to 184 | 9.0 ± 0.2 | 1.3x10<sup>2</sup> ± 8.8 | 2.45±3.53 | N<sub>2</sub> & H<sub>2</sub> off/on |
| | 3 | 185 to 234 | 8.9 ± 0.3 | 1.4x10<sup>2</sup> ± 8.2 | 6.14±7.00 | N<sub>2</sub>, H<sub>2</sub> |
| R3 | 1 | 75 to 142 | 7.8 ± 0.3 | 1.6x10<sup>2</sup> ± 7.6 | 0.50±0.31 | No gas |
| | 2 | 143 to 184 | 8.4 ± 0.5 | 1.3x10<sup>2</sup> ± 13 | 5.02±5.64 | N<sub>2</sub> off/on |
| | 3 | 185 to 234 | 7.0 ± 0.6 | 1.8x10<sup>2</sup> ± 15 | 10.2±4.83 | N<sub>2</sub> |

Our results show that the H_2_ partial pressure is a sensitive parameter to the *n*-caprylate performance, amplifying minor differences in operating conditions. During Period 1, gas in the headspace of Reactor 3 contained 31 ± 9.6% H_2_ (by volume), whereas H_2_ was 9.9 ± 5.2% and 1.8 ± 1.9% of total gas for Reactors 1 and 2, respectively (**Table S1**). H_2_ is produced *via* the RBOX pathway (**Fig. 1**). The reactor tightness and material diffusiveness may influence the H_2_ partial pressures because H_2_, as the smallest molecule, may easily diffuse out of the system, while other gases would not. We built almost the entire reactor setup out of stainless steel to minimize H_2_ diffusion through plastic tubing and connections. However, our results show that we were not able to prevent H_2_ diffusion out of the system, which included a gas recirculation pump and some tubing lines that were not made of stainless steel.

The H_2_ partial pressure can negatively affect chain elongation by reducing chain-elongating rates and changing the product spectra (21, 37). High H_2_ partial pressures can lower the ethanol oxidation rate to acetate and prevent the Rnf complex from functioning properly within the RBOX pathway, resulting in lower ATP production through substrate-level phosphorylation and membrane-based phosphorylation, respectively (21). This would lower the growth rate, further slowing the development of an active microbiota. With the relatively high H_2_ partial pressures for Reactor 3 during Period 1 compared to Reactors 1 and 2, a significant fraction of ethanol that we fed to Reactor 3 was not converted and left in the effluent, which resulted in a higher average effluent ethanol concentration for Reactor 3 compared to Reactor 1 and 2 (1.7×10^2^ ± 9.7 mM *vs*. 47 ± 3.9 mM and 29 ± 4.3 mM, respectively (**Fig. S2**, **Table S2**). Because our reactors were continuously stirred systems, concentrations measured in the effluent were approximately equal to what the reactor microbiomes observed.

To test whether a lower H_2_ partial pressure would improve *n*-caprylate production rates, we sparged N_2_ gas into Reactor 3 to reduce the percentage of H_2_ in the headspace (**Table 1; Table S1**). The sparging decreased the H_2_ in the headspace from 31 ± 9.6% (by volume) during Period 1 to 20 ± 14 % during Period 2 to 7.3 ± 4.6% during Period 3 (**Table S1**), resulting in increased volumetric *n*-caprylate production rates for Reactor 3 during Periods 2 and 3 (**Fig. 2B**). Into Reactor 2, we sparged N_2_ and H_2_ gas into the reactor. As expected, when hydrogen partial pressures increased during Periods 2 and 3 (**Table S1)**, *n*-caprylate productivity decreased for Reactor 2 (**Figure 2B**). However, we observed that the effect of H_2_ on *n*-caprylate production was not uniform in all reactors. When the amount of H_2_ in the headspace decreased due to N_2_ sparging into Reactor 1, we observed decreased *n*-caprylate production rates during Periods 2 and 3 (**Fig. 2B; Table S1**). However, sparing with N_2_ to remove H_2_ may have also removed O_2_, which could have an unknown effect. Gas sparging itself was another introduced variable in the experiment that may have decreased biomass growth and *n*-caprylate production for Reactor 1 during Periods 2 and 3. We also noted differences in the acetate, *n*-butyrate, *n*-caproate, and *n*-caprylate concentrations in the effluent of our reactors (**Table S2, Fig. S2A-C**). Thus, our system was not predictive because we did not fully understand how the environmental conditions in the reactor affect the microbial pathways in the complex microbiota.

### Bacterial species abundance correlated with *n*-caprylate production rates

We analyzed the reactor microbiome *via* 16S rRNA gene sequencing and shotgun metagenomic sequencing. Overall, we observed similar trends in the dominance of certain bacterial species during high and low *n*-caprylate production periods in both datasets. We noticed some differences between the sequencing methods, which we attributed to differences in how the data was analyzed and how taxonomy was assigned (see Methods). The 16S rRNA gene sequencing dataset was derived from approximately weekly biomass samples, which we collected from the reactors throughout the operating period. The shotgun metagenomics dataset was smaller and was derived from nine biomass samples, which we collected from the three reactors at three time points during the operating period (during Periods 1D, 3B, and 3C, as indicated in **Fig. 3**). The shotgun metagenomic dataset resulted in 477,902,544 reads. We assembled 23 high-quality draft genomes from this data, as detailed in **Table 2**. For metagenome-assembled genomes (MAGs) with high contamination, we observed multiple instances of single-copy genes, likely from the same or closely related strains, indicating contamination.

**Figure 3.**
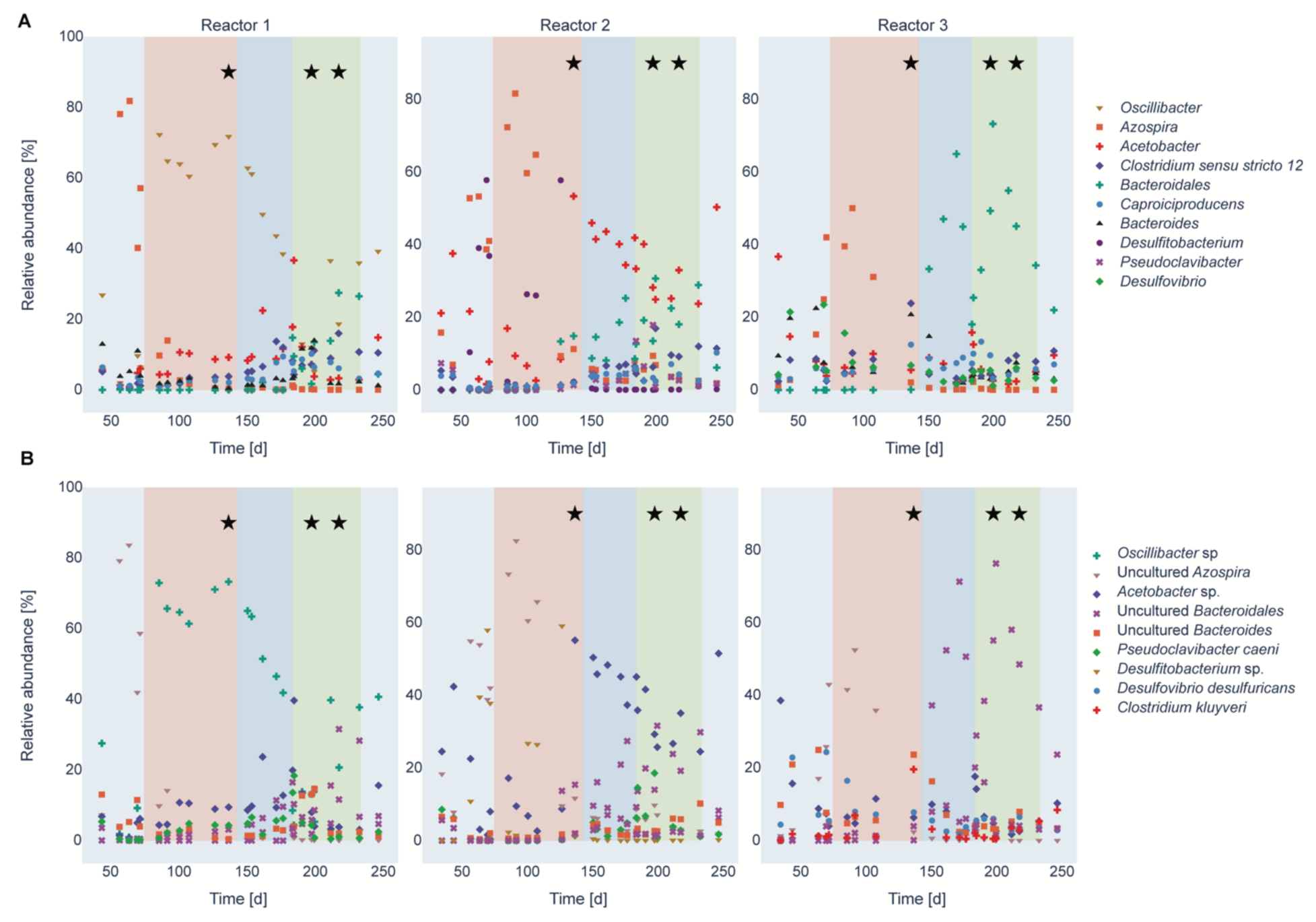
Relative abundance of the top seven most dominant taxa of each reactor based on the Illumina 16S rRNA gene sequencing results on the genus level (**A**) and the species level (**B**) throughout the operating time. The first 75 days of the operating period were the startup period (light blue). The salmon, blue, and green shadings indicate Periods 1, 2, and 3, respectively; the stars indicate the metagenomic sampling time points.

**Table 2.** Complete (>85%) metagenome-assembled genomes (MAGs) found in the reactors. The table depicts the assigned taxonomy of the bins, the completeness and contamination computed by CheckM, and the corresponding period in the reactor. If a MAG was present in multiple reactors, we only depicted the one with the highest completeness. The table is sorted by completeness.

| MAG | Completeness (%) | Contamination (%) | Period |
| --- | --- | --- | --- |
| *Methanobrevibacter* | 99.2 | 0.86 | 1D |
| *Prevotella paludivivens* | 98.7 | 0.79 | 3B |
| *Pseudoclavibacter caeni* | 98.3 | 0.58 | 1D |
| *Lactobacillus brevis* | 96.9 | 0 | 3C |
| *Clostridium* | 96.5 | 74.7 | 3C |
| *Acetobacter* | 95.2 | 34.7 | 1D |
| *Lactobacillus fuchuensis* | 94.47 | 2.36 | 3C |
| *Clostridium fermenticellae* JN500901 | 92.3 | 2.07 | 3C |
| *Lactobacillus sakei* | 91.9 | 1.09 | 1D |
| Bacteroidaceae | 91.4 | 35.3 | 1D |
| *Clostridium amylolyticum* | 88.2 | 6.50 | 1D |
| *Cellulomonas citrea* Ao-9 | 87.7 | 0.14 | 3B |
| *Oscillibacter* | 87.5 | 69.4 | 3B |

For both the 16S rRNA gene sequencing data and the shotgun metagenomics data, certain bacterial species within the genus *Oscillibacter* dominated when *n*-caprylate production rates were higher for Reactor 1 during Period 1 and decreased in abundance in later periods when production rates decreased (**Fig. 3B**, **Fig. 4**). The unknown *Oscillibacter* sp. bacteria that was dominant in the 16S rRNA gene sequencing data had a 95.9% ID to an *Oscillibacter valericigenes* Sjm18-20 strain (38). In the shotgun metagenomics analysis, *Oscillibacter valericigenes* was one of the dominant bacteria in Reactor 1 during Period 1 (126,083 aligned reads, **Fig. 4A**). Based on the shotgun metagenomics analysis, *O. valericigenes* abundance positively correlated with *n*-caprylate production rates (Pearson correlation coefficient, r=0.68, p=0.0439). Several other *Oscillibacter* spp. were also positively correlated with *n*-caprylate production rates, *Oscillibacte*r sp. CAG:155 (r=0.71, p=0.0321), *Oscillibacter ruminantium* (r=0.65, p=0.058), *Oscillibacter* sp. 1-3 (r=0.72, p=0.0287), *Oscillibacter* sp. NSJ-62 (r=0.65, p=0.058), and *Oscillibacter* sp. PC13 (r=0.68, p=0.0439) (**Fig. 4**). Based on the 16S rRNA gene sequencing data, an *Oscillibacter* sp. OTU, an uncultured *Oscillibacter* OTU, and an *Oscillibacter valericigenes* OTU were positively correlated with *n*-caprylate production rates (r=0.38, 0.27, and 0.15, and p=0.001, 0.021, 0.208 respectively, **Fig. 3**).

**Figure 4.**
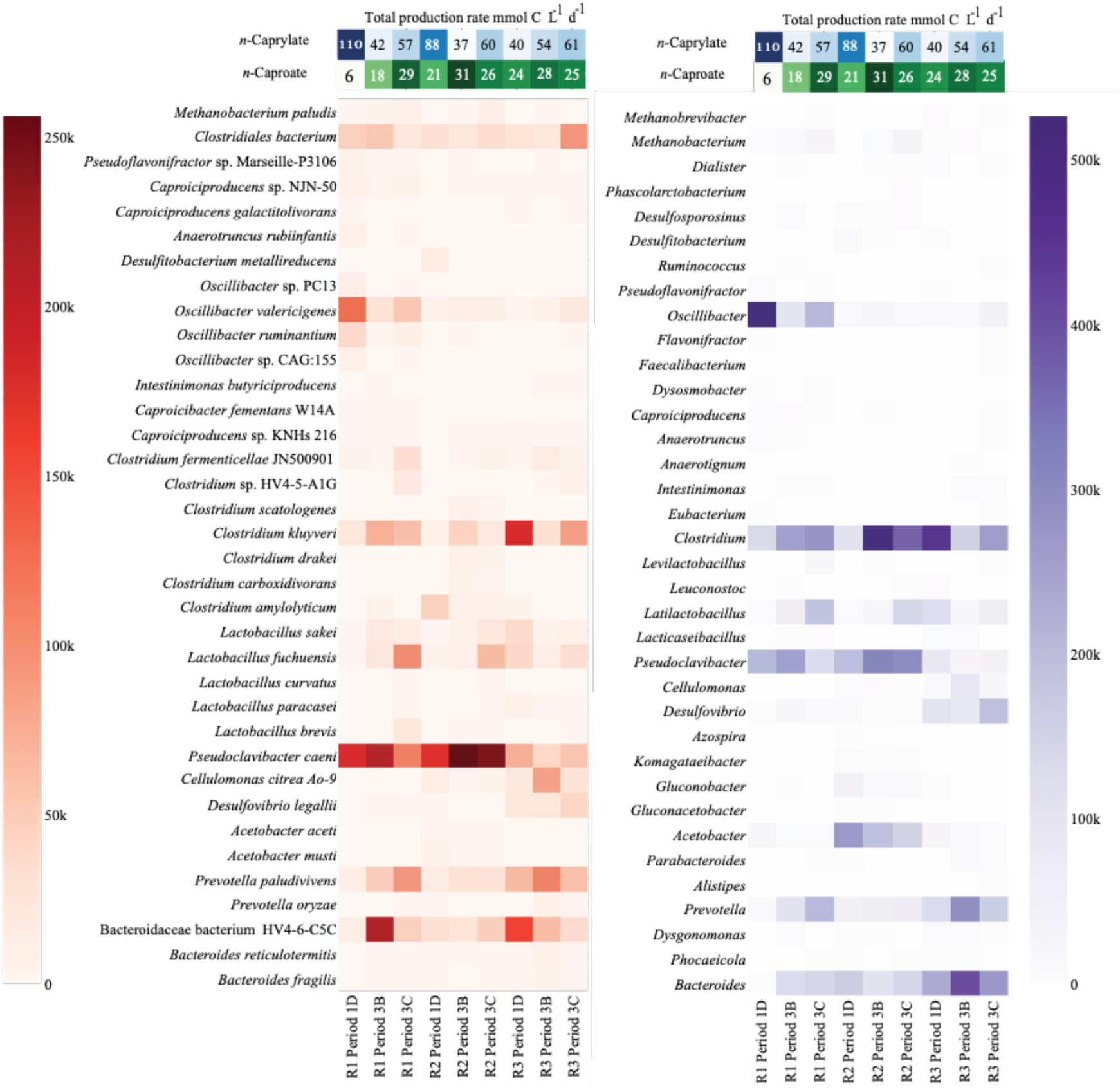
The most abundant species (**A**) and genus (**B**) in the three reactors based on the shotgun metagenome analysis. After normalizing the read count for sample size, the heatmap shows the number of reads aligned to each taxon throughout different sampling points of the reactors. Only taxa with more than 12k reads aligned are displayed. The names are ordered based on the NCBI taxonomy. The top of the plot shows the *n*-caprylate (blue) and *n*-caproate (green) volumetric production rates for Periods 1D, 3B, and 3C, respectively; color intensity is proportional to the production rates. R1-3 are Reactors 1-3.

Two aerobic bacteria, *Pseudoclavibacter caeni* and an unknown *Acetobacter* sp., were present in some reactors at relatively high abundances. For Reactors 1 and 2, *P. caeni* was an abundant bacterium. Still, its abundance did not correlate to *n*-caprylate production rates (r=0.01, p=0.98, **Fig. 4**). For reactor 2, *Acetobacter* sp. bacteria were dominant during Periods 1 and 2 and declined in abundance during Period 3 when *n*-caprylate production rates decreased (**Fig. 3 and 4**). The presence of these bacteria shows that O_2_ was introduced into the reactors due to an unknown location in the reactor setup.

Certain bacterial species dominated the reactors during periods of relatively low *n*-caprylate production but higher *n*-caproate production rates (Period 3B for Reactors 1 and 2 and Period 1D for Reactor 3; **Fig. 2**, **Fig. 3**, **Fig. 4**). Based on the shotgun metagenomics data, the abundance of *C. kluyveri* was negatively correlated to *n*-caprylate production rates, though the correlation was not significant (**Fig. 4**, r = -0.49, p=0.18). No correlation was observed between *C. kluyveri* relative abundance and production rates in the 16S rRNA gene sequencing data (**Fig. 3B**). For Reactor 1 during Period 3B, Bacteroidaceae bacterium HV4-6-C5C (217,314 aligned reads) and *P. caeni* (210,785 aligned reads) dominated the reactor. For Reactor 2 during Period 3B, *P. caeni* (255,949 aligned reads) and, to a lesser extent, *C. kluyveri* (43,228 aligned reads) dominated the reactor. For Reactor 3 during Period 1D, *C. kluyveri* (180,084 reads) and Bacteroidaceae bacterium HV4-6-C5C (157,401 reads) dominated the reactor (**Fig. 4**). Across all reactors, the abundance of Bacteroidaceae bacterium HV4-6-C5C was negatively correlated to *n*-caprylate production rates, though the correlation was not significant (r = -0.54, p= 0.133 **Fig. 4**). Based on the 16S rRNA gene sequencing data, an *Azospira* sp. was dominant in all the reactors *prior* to and at the start of Period 1 and was positively correlated to *n*-caproate production rates (r=0.41, p= 0.00034 **Fig 3**).

### Bacteria with the RBOX pathway

We investigated which metagenomes in our reactors had a complete or nearly complete RBOX pathway (**Fig. 5**). Specifically, we looked for nine enzymes involved in the RBOX pathway in our metagenomics and proteomics data: acetate CoA-transferase (CoAT), 3-hydroxy-acyl-CoA dehydrogenase (HAD), enoyl-CoA dehydratase (ECH), acyl-CoA dehydrogenase (ACD), electron-transfer-flavoprotein subunit A/B (EtfA/B), acetyl-coenzyme A acetyltransferase (ACAT), thioesterase (TE), and Rnf respiratory complex (RNF) (**Fig. 1**).

**Figure 5:**
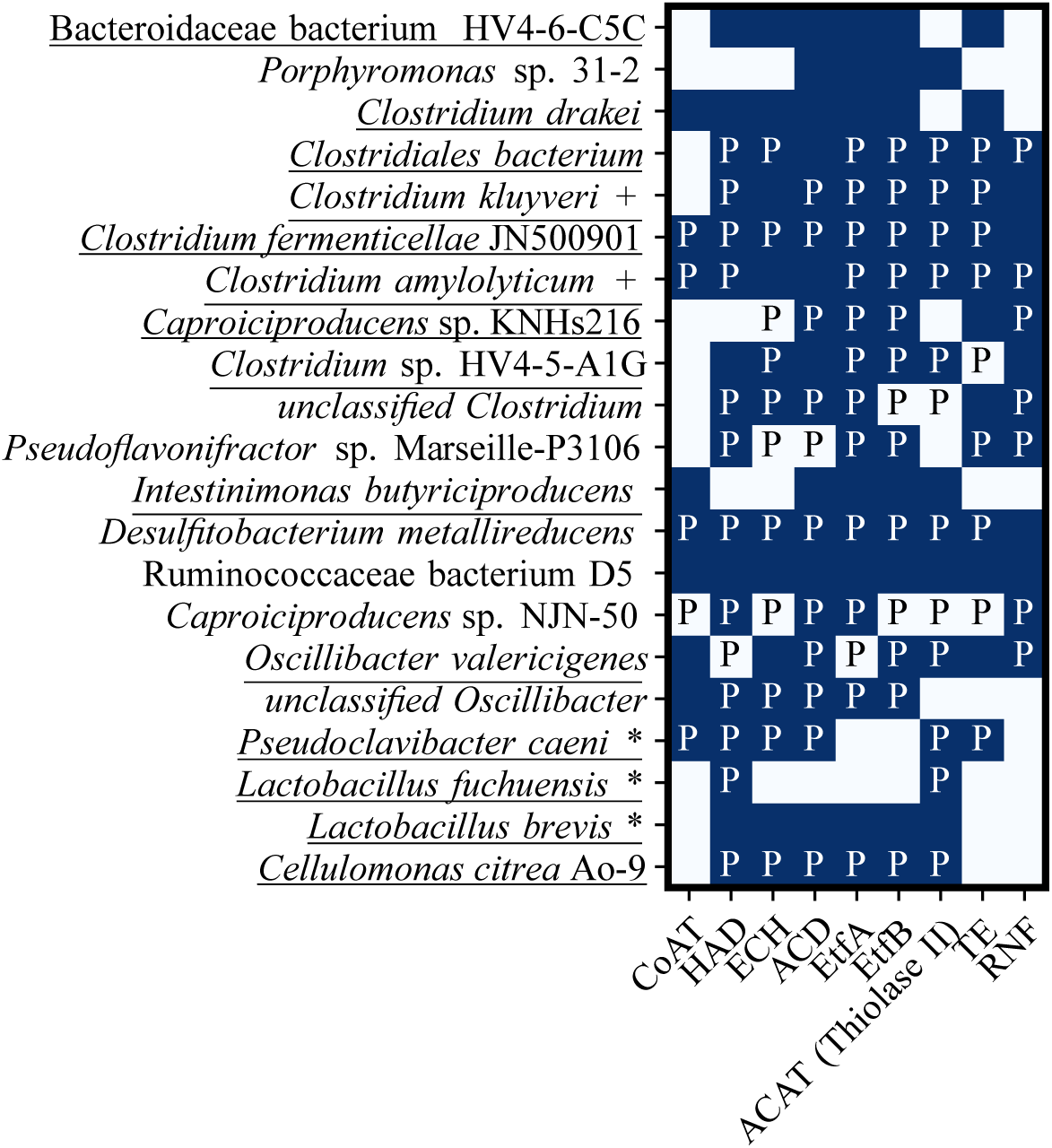
Absence or presence of nine enzymes involved in the RBOX pathway (acronyms described previously) in reactor *de-novo* assembled metagenomes and proteomes as monitored by shotgun metagenomics analysis and proteomics. Enzyme acronyms were described in **Fig. 1**. A blue box denotes the presence in the metagenome, the letter P the presence in the metaproteome, and a white box without a P the absence in both the metagenome and metaproteome. MAGs identified to species level are depicted. A * indicates >90% complete and <5% contaminated (determined with CheckM). A + indicates >80% complete and <10% contamination. More abundant species above the 12k read threshold are underlined.

All bacteria with the complete RBOX pathway in their metagenome and proteome were in the class *Clostridia* (**Fig. 5**). The *Clostridia* class member *O. valericigenes*, which dominated Reactor 1 during periods of high *n*-caprylate production (**Fig. 3**), contained the majority of RBOX enzymes (except for enoyl-CoA dehydratase) in its metagenome and proteome (**Fig. 5**). Electron-transfer-flavoprotein subunit B was only detected in the proteomics data. Other bacteria in the class *Clostridia* had the complete RBOX pathway but were not dominant bacteria. *Clostridium fermenticellae* JN500901 and *Clostridium amylolyticum* had complete RBOX pathways in their metagenomes, though, the pathway was not complete in the proteomics data (**Fig. 5**). It is important to note that the absence of a protein in our comparative proteomics does not mean that the protein is not present. *Clostridium fermenticellae* JN500901 was detected in all reactor samples except for Reactor 2 Period 1D and was most abundant for Reactor 1 during Period 3C (32,522 aligned reads; **Fig. 4**). *Clostridium amylolyticum* was most abundant for Reactor 2 during Period 1D (45,149 aligned reads; **Fig. 4**). *Clostridium* sp. HV4-5-A1G, which was most abundant for Reactor 1 during Period 3C (18,954 aligned reads; **Fig. 4**), had a nearly complete RBOX pathway and was only missing genes for acetate CoA-transferase enzyme (**Fig. 5**).

We found the complete RBOX pathway in the proteomics but not the metagenomics data for *Caproiciproducens* sp. NJN-50 (**Fig. 5**), which was present in all metagenomics timepoints except for Reactor 2 during Period 1D. In a *prior* study, which utilized inoculum from this study’s reactors, another *Caproiciproducens* strain (7D4C2) was isolated from reactor biomass and shown to produce *n*-caproate (39). *Desulfitobacterium metallireducens* contained the entire RBOX pathway in its genome and proteome (except for RNF respiratory complex, which was only found in the genome) (**Fig. 5**). This bacterium was only found for Reactor 2 Period 1D (16,507 aligned reads, **Fig. 4**). Ruminococcaceae bacterium D5, which was only found for Reactor 3 during Periods 3B and 3C (4201 and 3544 aligned reads, respectively; **Fig. 4**) had the complete RBOX pathway in its metagenome but not in its proteome (**Fig. 5**). *C. kluyveri* and *Clostridiales* bacterium metagenomes contained and expressed the majority of RBOX enzymes except for acetate CoA-transferase (**Fig. 5**).

### Bacterial microcompartments present in reactor microbiomes

The metagenomic analysis revealed the presence of specific bacterial microcompartments. These specialized compartments encapsulate metabolic pathways, enhancing metabolic efficiency and specificity. It is known that the chain-elongator *C. kluyveri* and other clostridia have microcompartments to protect its intracellular milieu against unstable or toxic chemical intermediates (40). Notably, an ethanolamine utilizing microcompartment (EUT2B), two propanediol utilizing microcompartments (PDU1D and PDU1C), and a glycyl radical enzyme containing microcompartment (GRM1A) were the dominant microcompartments observed (**Fig. S3**). These microcompartments are metabolosomes that are usually expressed only when their substrate is present (41). The EUT microcompartment metabolizes ethanolamine, which is a product of the breakdown of phosphatidylethanolamine, to ethanol, acetyl-CoA, and acetyl-phosphate and protects the rest of the bacterial cell from the intermediate acetaldehyde (42). We also observed the presence of the ethanol utilizing microcompartment (ETU) in all time points studied (**Fig. S3**). This bacterial microcompartment has only been reported in the bacterium *C. kluyveri* (43, 44). In our analysis, we observed the ETU microcompartment as expected in *C. kluyveri*. We also observed the ETU microcompartment in other bacteria classified to the level *Clostridium* spp. or Clostridiaceae (**Fig. S3**), though, we note that these could be *C. kluyveri* species that cannot be classified to the species level. We also observed the ETU microcompartment in proteins that had no hit in the taxonomic database (NAs in **Fig. S3**). *Oscillibacter valericigenes* Sjm18-20, which dominated Reactor 1 during periods of high *n*-caprylate production, was not found to have ETU microcompartments, though, it did contain EUT2B, PDU1D, and GRM1A, GRM3A, and GRM3C microcompartments.

## Discussion

For our open-culture reactors, different microbial communities were correlated with periods of high *n*-caproate or *n*-caprylate production (**Fig. 4**). The known chain elongator *C. kluyveri* and a primary fermenter Bacteroidaceae bacterium HV4-6-C5C had higher relative abundances during periods of high *n*-caproate production and decreased in abundance during periods of high *n*-caprylate production (**Fig. 3-4**). *C. kluyveri* produces *n*-caproate from ethanol and short-chain carboxylates (acetate or *n*-butyrate), but there is limited evidence of its ability to produce *n*-caprylate (10). In particular, the ethanol-utilizing microcompartment, ETU, associated with *C. kluyveri*, underscores its potential role in the efficient conversion of ethanol to acetaldehyde, which is a pivotal step in the production of *n*-caproate and *n*-caprylate. Different *Oscillibacter* species, which include *Oscillibacter valericigenes* (r=0.68, p=0.0439), were positively correlated to periods of high *n*-caprylate output in the reactors (**Fig. 3-4**). Indeed, *O. valericigenes* included microcompartments to possibly protect itself during chain elongation. This finding is consistent with *prior* studies for which members of the Ruminococcaceae family (to which *Oscillibacter* belongs) were isolated from reactors producing *n*-caproate from lactate (9, 29) and Illumina 16S rRNA gene sequencing studies for which Ruminococcaceae members were associated with medium-chain carboxylate production in reactors (4, 7, 11).

The unplanned presence of O_2_ in our reactors created a niche for aerobic bacteria, such as *P. caeni* and *Acetobacter* species, to survive and become abundant in the reactors (**Fig. 4)**. The abundance of these aerobic bacteria was not correlated to *n*-caprylate production rates (**Figs. 3-4**). As a result of our inability to build a reactor system that prevented O_2_ inclusion, a major caveat existed in our quest to study different H_2_ partial pressures on the RBOX. The use of gas sparging to remove or add H_2_, also removed O_2_, and this turned out to be a sensitive parameter. Even though, we could not satisfy our experimental design with the independent parameter H_2_, this study is providing us with information to base future research on, as discussed below. Aerobic or facultative anaerobic microbes must have quickly consumed the O_2_ in our reactors because strict anaerobic microbes, such as methanogens and other obligate anaerobes, were also present in our continuously stirred reactor systems (**Fig. 4**). *Prior* studies observed aerobes, such as *Acetobacter* (3, 11), and facultative anaerobes, such as *Lactobacillus* (11, 45), in chain elongation reactors. Previous studies from our lab had not found *P. caeni* in similar chain-elongating reactors (3, 4), though, the aerobe *Acetobacter* was observed (3). *P. caeni* was isolated from sewage sludge in 2012 (46), but the *P. caeni* assembly was only added to the NCBI nr database in 2019 (ASM883112v1). *P. caeni* could have been present in previous reactor studies but not detected due to its absence from existing databases. A previous study from 2016 found a phylotype that matches *P. caeni* in batch experiments utilizing biomass from a chain-elongating reactor that was fed a variety of substrates and found its occurrence did not correspond to chain elongation activity (47).

From our metagenomic and metaproteomic analyses, we conclude that the reverse *β*-oxidation pathway was active in our reactors (**Fig. 5**). Some abundant bacteria did not have the complete RBOX pathway (**Fig. 5**), which may indicate: **(1)** that our methods did not always identify all genes or proteins; or **(2)** that chain elongators live in syntropy with each other to produce the medium-chain carboxylic acids. We observed that RBOX was affected by the partial pressures of H_2_ in the headspace of the reactors, which follows the current understanding of chain elongation. The presence and distribution of specific bacterial microcompartments in the dominant bacteria (**Fig. S3**) could influence this observed metabolic pathway, reflecting the potential versatility introduced by these microcompartments. We should note that we only have evidence that genes for the bacterial microcompartments were present in the genomes, not that they were expressed. As expected, *C. kluyveri* had most of the RBOX pathway in the metagenome, the only exception being CoAT. We did not find RNF and ECH proteins in the proteome for this bacterium but only in the metagenome. The *O. valericigenes* species bacterium, which was dominant during the period of highest *n*-caprylate production, had the nearly complete RBOX pathway. Other *Clostridium* species, *Clostridium fermenticellae* JN500901*, C. amylolyticum, Desulfitobacterium metallireducens,* Ruminococcaceae bacterium D5, and *Caproiciproducens* sp. NJN-50 had a complete or nearly complete RBOX pathway (**Fig. 5**). In general, these were not the dominant bacteria in our reactors (**Fig. 4**). We also note that some bacteria found in our study, including the aerobe *P. caeni*, had the complete or nearly complete RBOX pathway but are not known chain elongators. Previous researchers have also noted the presence of the RBOX pathway in bacteria that are not known chain elongators (39).

Our study provides insight into the bacteria producing *n*-caproate and *n*-caprylate from ethanol and acetate *via* reverse *β*-oxidation. We identified potential candidates for *n*-caprylate production in our reactors. Future studies should try to isolate and sequence *n*-caprylate-producing bacteria. In addition, future studies should also investigate whether a relative lack of diversity in *n*-caprylate-producing reactors affects the stability of these systems. The potential influence of bacterial microcompartments on this metabolic pathway, as observed in our shotgun metagenomic analysis, underscores the need to consider their role in future studies. Future research should also investigate the role of microaerobic conditions in these reactors because we observed that O_2_ is a sensitive parameter, but we do not know why.

## Materials and Methods

### Continuously Fed Reactor System

We designed, self-built, and operated three grade-316 stainless steel reactors with a 5.5-L total volume (5-L working volume and 0.5-L headspace volume) (parts utilized in the building system detailed in **Table S3**). We maintained the reactor pH at ∼5.5 (*via* periodic additions of 0.5 M HCl) and the temperature at 30 ± 1.0°C. We fed the reactors with a modified-based media that was previously described (4, 48) and supplemented with ethanol and acetate. After a 75-day startup period, we mixed broth from all reactors to ensure similar microbiota in each reactor before a restart. We operated the reactors as replicates in which we kept organic loading rates and hydraulic retention time at 1.5×10^2^ ± 4.6 mM C L^-1^ d^-1^ and 8.5 ± 0.2 days, respectively, for a period of 68 days (Period 1 of study – Days 75 to 142; see **Table 1**).

Reactors were inoculated with 10% by volume (∼500 mL) of reactor broth from a reactor that was fed with ethanol-rich yeast fermentation beer and operated as an anaerobic sequencing batch reactor for an operating period of approximately five years *prior* to the time we collected the inoculum (7, 16). For in-line product extraction, we used a setup such as the one previously described by Agler et al. (7). Detailed information on the reactor setup can be found in the Supporting Information. During Periods 2 and 3, gases (N_2_ and H_2_) were sparged into the bottom port of the reactors (**Table 1)**.

### Experimental Periods for Reactors

The primary study periods were Periods 1 (Days 75 to 142), Period 2 (Days 143 to 184), and Period 3 (Days 185 to 234), and were divided into periods: 1A – Days 75 to 91, 1B – Days 92 to 113, 1C – Days 114 to 125, 1D – Days 126 to 142, 2A – Days 143 to 151, 2B – Days 152 to 162, 2C – Days 163 to 184), 3A – Days 185 to 194, 3B – Days 195 to 206, and 3C – Days 207 to 222 (**Table 1**). *Prior* to Period 1, there was a 75-day startup period for the reactors in which the organic loading rate was incrementally increased to the target loading rate of ∼140 mM C L^-1^ d^-1^ at a hydraulic retention time of ∼9 days. At the start of Period 1 (Day 75), biomass from all three reactors was combined, mixed, and redistributed. During Period 1, operating conditions (*i.e.*, temperature, pH, product extraction) were kept the same between the three reactors except for minor differences in organic loading rates that we applied. During Period 2, gas sparging of N_2_ gas was tested out (*i.e.*, gas sparging was off and on between Days 143 to 184) (**Table 1**). At the start of Period 3 (Day 185), biomass was again mixed and redistributed. During Period 3, we sparged Reactor 1 and Reactor 3 continuously with N_2_ gas, while we sparged Reactor 2 continuously with a mixture of H_2_ and N_2_ gas (**Table 1)**. Although we did not measure the gas flow rate that we sparged into the reactors during Period 3, sparging rates are assumed to be equal to the measured exit gas flow rates reported due to low gas production rates that we observed during Period 1 without sparging (**Table 1**).

### Liquid and Gas Analysis

We collected liquid samples from reactor broth and alkaline extraction solution to measure carboxylate and ethanol concentrations. The 2-mL samples of reactor broth were collected from a port in the system recycle line of the reactor. In contrast, we collected the alkaline extraction solution samples from a ∼3-L well-mixed glass reservoir from which the extraction solution was re-circulated. Samples were stored frozen at -20°C *prior* to analysis. Gas chromatography systems were used to determine carboxylate and ethanol concentrations, as has been described by Usack et al. (49). We collected gas samples from gas exit lines of the reactors. CO_2_, CH_4_, and H_2_ concentrations (>0.2% by volume) were measured using a gas chromatography system, which has been described previously (49). A reduction gas detector (RGD) was used to measure H_2_ gas concentrations <0.2%, which has been described by Kucek et al. (4).

### Calculations and Statistical Analysis of Operating Data

We calculated the carboxylate production rates as average values for each operating period. We summed the average effluent production rates per liter of the reactor (mmol C L^-1^ d^-1^) and the average transfer rates *via* product extraction (mmol C L^-1^ d^-1^) to yield total production rates per liter of the reactor (mmol C L^-1^ d^-1^). We calculated the average effluent production rates by dividing the average carboxylate concentration per period by the average HRT. We calculated the average HRT per period based on the average effluent flow rate per period, which was determined gravimetrically. We calculated the average transfer rates by plotting the increasing concentrations of individual carboxylates in alkaline extraction solution *vs.* time. We used least squares methods to determine the slope and the sample standard deviation (LINEST function, Microsoft Excel). We divided the slope by the reactor working volume (5 L) to obtain an average transfer rate per period. RStudio v.1.0.136 (50) was used to run data analysis in R. Concentrations, rates, ratios, and efficiencies are reported as mean value ± standard error in the paper unless noted otherwise.

### *16S* rRNA Gene Sequencing Analysis

We took close-to-weekly biomass samples for Illumina 16S rRNA gene sequencing analysis from the internal recycle line of the reactors. Approximately 10 mL of reactor broth was collected with a 60-mL plastic syringe and distributed into 2-mL Eppendorf tubes. We centrifuged the tubes at 16,873 x *g* for four min and discarded the supernatant. Finally, we stored the pelleted biomass samples at -80°C.

According to the manufacturer’s protocol, we extracted genomic DNA using the PowerSoil-htp 96 Well Soil DNA Isolation kit (MO BIO Laboratories Inc., Carlsbad, CA). DNA amplification protocol was described previously (51) with the following exceptions: Mag-Bind RxnPure Plus magnetic beads solution (Omega Biotek, Norcross, GA, USA) was used instead of Mag-Bind E-Z Pure, and 50 ng DNA per sample was pooled instead of 100 ng. Duplicate PCR reactions of each DNA extract were performed and pooled *prior* to sequencing. Samples were sent for paired-end sequencing (2×250bp) on the Illumina MiSeq platform (Illumina, San Diego, CA, USA) at the Cornell University Biotechnology Resource Center (Ithaca, NY, USA). We analyzed the resulting 16S rRNA gene sequencing reads using QIIME 2 2017.3 (52). Finally, we investigated the correlation of the relative abundance for OTUs with *n*-caproate and *n*-caprylate production rates using the scipy-stats package pearsonnr (53).

### Shotgun Metagenomic analysis

We collected biomass samples for shotgun metagenomic analysis from internal liquid-recycle lines of the reactors, which were utilized to mix the reactor liquid. We obtained three samples for each reactor: during Period 1C on Day 137 for Reactors 1 and 2 and a pooled sample for Days 137, 151, 154, and 162 for Reactor 3, Period 3B on Days 198 and 200, and Period 3C on Day 218). Samples were centrifuged, supernatant was discarded, and biomass was stored at -80°C. Genomic DNA was extracted using the PowerSoil DNA Isolation kit (MO BIO Laboratories Inc., Carlsbad, CA). We used a modified protocol, which has been described by Kucek et al. (4). Nine extracted DNA samples were barcoded and sequenced on two lanes (100 bp *per* read; single-direction reads) using Illumina HiSeq platform at the JP Sulzberger Genome Center at Columbia University (New York, New York). We merged the replicates of samples.

Shotgun metagenomics read quality was checked using FastQC (54) after trimming with Trimmomatic (55). We performed quality control using FastQC (54) version 0.11.9 on merged reads. The sequence quality scores and histograms passed standard test criteria for all samples (lower quartile for every base above 10 and median above 25). To trim low-quality regions and remove low-quality reads, Trimmomatic (55) version 0.39. was applied on all samples providing the parameters -phred33; LEADING:3; TRAILING:3 as well as SLIDINGWINDOW::4:15 and MINLEN:36. Trimmed reads were aligned to NCBI-nr database (Feb 2021) using DIAMOND (56) version 2.0.7.in blastx mode. The following parameters were used: --outfmt 100 -c1 -b12 -p 32 --top 10 -e 0.001. Resulting alignments were meganized for further analysis using daa-meganizer, which is a tool that is included in MEGAN6 (57). DIAMOND output files were loaded into MEGAN6, were normalized by sample size, and read counts were extracted for each MAG. Heatmaps were created using a Python script, only displaying MAGs with more than 12k aligned reads (**Fig. 4**). The correlation of read counts with *n*-caproate and *n*-caprylate production rates was investigated using Pearson correlation coefficient (53).

### De-novo *Assembly*

We performed de novo assembly for each set of quality filtered reads using MEGAHIT (58) version 1.2.9 with preset -meta-large for large and complex metagenomes. This resulted in 757,643 contigs with a mean length of 1129 bp and a mean N50 of 2620 bp. Assembled contigs were aligned to NCBI-nr using DIAMOND blastx mode and parameters –long reads --outfmt 100 -c1 -b12. The resulting files were meganized using daa-meganizer, using the parameters - mpi 80 -top 10. We extracted the binned contigs using MEGAN’s read-extractor tool, resulting in one fasta file per bin. On average, 214 bins were created per sample. The bins were checked for completeness and contamination using CheckM (59) version 1.1.2 in lineage_wf mode. We annotated the contigs using Prokka (60) version 1.13.3. We created a custom HMM (hidden Markov model) to search for the absence and presence of genes involved in the RBOX pathway. We based the included genes on the models used by Scarborough et al. (25). Pre-trained HMMs were downloaded from PFAM (61) and all annotated proteins from each sample were searched with these models using HMMER3 (62). For the RBOX pathway, PFAM only has 3-hydroxyacyl-CoA dehydrogenase (HAD) and not 3-hydroxy-butyrl-CoA dehydrogenase (HBD), so HBD was omitted. A custom Python script was used to plot the presence or absence of each gene in the annotated proteins. The annotated proteins were also searched for biological microcompartment (BMC) proteins using the BMC Caller tool (63) and plotted with Python (**Fig. S3**).

The samples from Periods 1D and 3C for Reactor 1 failed the per-base-sequence-content test, while samples from: **(1)** Periods 1D and 3C for Reactor 1; **(2)** Period 1C for Reactor 2; and **(3)** Period 3B for Reactor 3 all failed the per-sequence-GC-content test. Sequence duplication level was high in all samples except for the sample from Period 3C from Reactor 2. This problem was introduced when merging two replicates for each sample and is an artifact. Overall, the per-sequence quality scores were sufficient.

### Metaproteomic Analysis

For metaproteomic sampling, approximately 200 mL of reactor broth was collected from the internal recycle lines of the reactors and distributed into four 50 mL centrifuge tubes. After centrifugation for 10 min at 8000 x *g* (at 4°C), the supernatant was discarded. Pellets were resuspended in a tris buffer solution and redistributed to 2-mL Eppendorf tubes. We spun the tubes for 4 min at 16873 x *g* and discarded the supernatant. Next, we stored the pelleted samples at -20°C.

Protein samples were extracted from reactor cell pellets (∼100 μL bulk volume) using a gel-free, precipitation-free method to avoid loss of hydrophobic proteins. Cell pellets were suspended in 500 μL 50 mM Tris buffer (pH 8.0) and flash frozen 3 x with liquid N_2_ as an initial lysis step. 0.1% SDS, 10 mM NaCl, 0.02 M TCEP, and 2 M urea were added to lyse the sample by ultrasonication on ice at 60% amplitude for 5 min total pulse time, vortexed, and centrifuged 10 min at 12,000 x *g*. Half of the supernatant (∼250 μL) was removed and saved. To attempt to desorb more hydrophobic proteins from the pellet, 250 μL of acetonitrile was added to the cell pellet and the remaining supernatant. This was then vortexed and pelleted, while supernatant from this step was removed and re-combined with the first 250 μL of supernatant. The volume of the combined supernatant was decreased to approximately 400 μL *via* speed vac. We discarded the insoluble pellet. Total protein estimates were measured by Bradford assay. Protein samples were reduced with an additional 0.05 M TCEP in 0.1 M ammonium bicarbonate at 35°C for 1 h, alkylated with 40 mM iodoacetamide at room temperature for 30 min, and digested with Pierce Trypsin Protease MS-Grade at an estimated 1:20 trypsin: protein mass ratio for 12 h at 35°C with 1 mM CaCl_2_. Sample protein precipitation was avoided during digestion by diluting trypsin protease in 0.1 M ammonium bicarbonate buffer containing 0.02% SDS and 10% acetonitrile before combining with the protein sample. To quench digestion, samples were acidified to a pH of 3.5 with formic acid, acetonitrile was removed *via* speed-vac, acidified again to pH 3.5, and stored at -20°C. Tryptic peptides were purified using 1 mL Supelclean ENVI-18 SPE tubes and dissolved in 0.1% TFA / 0.5% acetonitrile for analysis by LC-MS.

LC-MS was performed using a Thermo Fisher UltiMate 3000 LC and LTQ-XL mass spectrometer with a standard ESI source. Microflow chromatography was performed on an Acclaim PepMap 100 column (1 mm x 15 cm; 3 um) at 40 μL/min using a 125 min gradient from 100 % water (1% formic acid) to 40 % acetonitrile. We operated the LTQ-XL in a 3x double play mode with a 10 s dynamic exclusion time and CID activation. The resulting peptides were compared to a decoy search. Peptides were thrown out based on a probabilistic filter. Proteins were kept if they had at least two unique peptides IDed with high confidence. The resulting 341 protein sequences were aligned against NCBI-nr (Feb 2021) using DIAMOND blastp version 2.0.7 for taxonomic assignment. To check for proteins involved in RBOX, we searched for these proteins using the previously described HMM models.

### Data Availability

16S rRNA gene sequences are available at EBI (https://www.ebi.ac.uk/) under accession number ERP024135. Sequences and study metadata are publicly available in QIITA (https://qiita.ucsd.edu/) under study number 11227. Shotgun metagenomics data is available at SRA (https://www.ncbi.nlm.nih.gov/sra) under the accession PRJNA824684. Metagenomics and metaproteomics data analysis code is available on GitHub (https://github.com/lucass122/caprylate_reactor_paper).

## Supporting information

r1t1_rbox_counts.csv

r2t1_rbox_counts.csv

BMC_from_BMC_caller_tool.csv

## Acknowledgments

The authors would like to acknowledge Chase Brett and Doug Caveney for their help with constructing the reactor and Alex Marzelli and Dr. Jiajie Xu for assistance with reactor maintenance (all from Cornell University). We acknowledge funding from the U.S. EPA STAR grant fellowship, the U.S. Army Research Laboratory, and the U.S. Army Research Office under contract/grant number W911NF-12-1-0555. We also acknowledge funding from the Alexander von Humboldt Foundation in the framework of the Alexander von Humboldt Professorship to LTA, the Novo Nordisk Foundation CO_2_ Research Center with grant number NNF21SA0072700 to LTA, the Deutsche Forschungsgemeinschaft under Germany’s Excellence Strategy (EXC 2124 – 390838134) to LTA and DH. A special thanks goes to the Reinhard Frank Stiftung to support the exchanges between the University of Maryland and the University of Tübingen.

## Notes

### Competing Interest Statement

The authors have declared no competing interest.

## References

1. Stamatopoulou P, Malkowski J, Conrado L, Brown K, Scarborough M. 2020. Fermentation of organic residues to beneficial chemicals: A review of medium-chain fatty acid production. Processes 8:1571.

2. Mancini A, Imperlini E, Nigro E, Montagnese C, Daniele A, Orrù S, Buono P. 2015. Biological and nutritional properties of palm oil and palmitic acid: Effects on health. Molecules 20:17339–17361.

3. Spirito CM, Marzilli AM, Angenent LT. 2018. Higher substrate ratios of ethanol to acetate steered chain elongation toward *n*-caprylate in a bioreactor with product extraction. Environmental Science & Technology 52:13438–13447.

4. Kucek LA, Spirito CM, Angenent LT. 2016. High *n*-caprylate productivities and specificities from dilute ethanol and acetate: Chain elongation with microbiomes to upgrade products from syngas fermentation. Energy & Environmental Science 9:3482–3494.

5. Grootscholten TIM, Steinbusch KJJ, Hamelers HVM, Buisman CJN. 2013. Chain elongation of acetate and ethanol in an upflow anaerobic filter for high rate MCFA production. Bioresource Technology 135:440–445.

6. Steinbusch KJ, Hamelers HVM, Plugge CM, Buisman CJN. 2011. Biological formation of *n*-caproate and *n*-caprylate from acetate: Fuel and chemicals from low grade biomass. Energy & Environmental Science 4:216–224.

7. Agler MT, Spirito CM, Usack JG, Werner JJ, Angenent LT. 2012. Chain elongation with reactor microbiomes: Upgrading dilute ethanol to medium-chain carboxylates. Energy & Environmental Science 5:8189–8192.

8. Kucek LA, Nguyen M, Angenent LT. 2016. Conversion of L-lactate into *n*-caproate by a continuously fed reactor microbiome. Water Research 93:163–171.

9. Zhu XY, Tao Y, Liang C, Li XZ, Wei N, Zhang WJ. 2015. The synthesis of *n*-caproate from lactate: A new efficient process for medium-chain carboxylates production. Scientific Reports 5:14360.

10. Gildemyn S, Molitor B, Usack JG, Nguyen M, Rabaey K, Angenent LT. 2017. Upgrading syngas fermentation effluent using *Clostridium kluyveri* in a continuous fermentation. Biotechnology for Biofuels 10:83.

11. Andersen SJ, De Groof V, Khor WC, Roume H, Props R, Coma M, Rabaey K. 2017. A *Clostridium* group IV species dominates and suppresses a mixed culture fermentation by tolerance to medium chain fatty acids products. Frontiers in Bioengineering and Biotechnology 5:8.

12. Carvajal-Arroyo JM, Candry P, Andersen SJ, Props R, Seviour T, Ganigué R, Rabaey K. 2019. Granular fermentation enables high rate caproic acid production from solid-free thin stillage. Green Chemistry 21:1330–1339.

13. Carvajal-Arroyo JM, Andersen SJ, Ganigué R, Rozendal RA, Angenent LT, Rabaey K. 2021. Production and extraction of medium chain carboxylic acids at a semi-pilot scale. Chemical Engineering Journal 416:127886.

14. Duber A, Jaroszynski L, Zagrodnik R, Chwialkowska J, Juzwa W, Ciesielski S, Oleskowicz-Popiel P. 2018. Exploiting the real wastewater potential for resource recovery – *n*-caproate production from acid whey. Green Chemistry 20:3790–3803.

15. Xu J, Guzman JJL, Andersen SJ, Rabaey K, Angenent LT. 2015. In-line and selective phase separation of medium-chain carboxylic acids using membrane electrolysis. Chemistry Communications 51:6847–6850.

16. Ge S, Usack JG, Spirito CM, Angenent LT. 2015. Long-term *n*-caproic acid production from yeast-fermentation beer in an anaerobic bioreactor with continuous product extraction. Environtal Science and Technology 49:8012-8021.

17. Khor WC, Andersen S, Vervaeren H, Rabaey K. 2017. Electricity-assisted production of caproic acid from grass. Biotechnology for Biofuels 10:180.

18. Grootscholten TIM, Kinsky dal Borgo F, Hamelers HVM, Buisman CJN. 2013. Promoting chain elongation in mixed culture acidification reactors by addition of ethanol. Biomass and Bioenergy 48:10-16.

19. Spirito CM, Richter H, Rabaey K, Stams AJM, Angenent LT. 2014. Chain elongation in anaerobic reactor microbiomes to recover resources from waste. Current Opinion in Biotechnology 27:115–122.

20. Cavalcante WdA, Leitão RC, Gehring TA, Angenent LT, Santaella ST. 2017. Anaerobic fermentation for *n*-caproic acid production: A review. Process Biochemistry 54:106–119.

21. Angenent LT, Richter H, Buckel W, Spirito CM, Steinbusch KJJ, Plugge CM, Strik DPBTB, Grootscholten TIM, Buisman CJN, Hamelers HVM. 2016. Chain elongation with reactor microbiomes: Open-culture biotechnology to produce biochemicals. Environmental Science & Technology 50:2796–2810.

22. Buckel W, Thauer RK. 2013. Energy conservation via electron bifurcating ferredoxin reduction and proton/Na+ translocating ferredoxin oxidation. Biochimica et Biophysica Acta (BBA) - Bioenergetics 1827:94–113.

23. Scarborough MJ, Hamilton JJ, Erb EA, Donohue TJ, Noguera DR. 2020. Diagnosing and predicting mixed-culture fermentations with unicellular and guild-based metabolic models. mSystems 5:e00755–00720.

24. Scarborough MJ, Myers KS, Donohue TJ, Noguera DR. 2020. Medium-chain fatty acid synthesis by “*Candidatus Weimeria bifida*” gen. Nov., sp. Nov., and “*Candidatus Pseudoramibacter fermentans*” sp. Nov. Applied and Environmental Microbiology 86:e02242–02219.

25. Scarborough MJ, Lawson CE, Hamilton JJ, Donohue TJ, Noguera DR. 2018. Metatranscriptomic and thermodynamic insights into medium-chain fatty acid production using an anaerobic microbiome. mSystems 3:e00221–00218.

26. Wu S-L, Sun J, Chen X, Wei W, Song L, Dai X, Ni B-J. 2020. Unveiling the mechanisms of medium-chain fatty acid production from waste activated sludge alkaline fermentation liquor through physiological, thermodynamic and metagenomic investigations. Water Research 169:115218.

27. Han W, He P, Shao L, Lü F. 2018. Metabolic interactions of a chain elongation microbiome. Applied and Environmental Microbiology 84:e01614–01618.

28. Gest H. 1995. A serendipic legacy: Erwin Eesmarch’s isolation of the first photosynthetic bacterium in pure culture. Photosynthetic Research 46:473–478.

29. Zhu X, Zhou Y, Wang Y, Wu T, Li X, Li D, Tao Y. 2017. Production of high-concentration *n*-caproic acid from lactate through fermentation using a newly isolated ruminococcaceae bacterium cpb6. Biotechnology for Biofuels 10:102.

30. Bornstein BT, Barker HA. 1948. The energy metabolism of *Clostridium kluyveri* and the synthesis of fatty acids. Journal of Biological Chemistry 172:659–669.

31. Barker HA, Kamen MD, Bornstein BT. 1945. The synthesis of *n*-butyric and *n*-caproic acids from ethanol and acetic acid by *Clostridium kluyveri*. Proceedings of the National Academy of Sciences 31:373–381.

32. Wallace RJ, McKain N, McEwan NR, Miyagawa E, Chaudhary LC, King TP, Walker ND, Apajalahti JHA, Newbold CJ. 2003. *Eubacterium pyruvativorans* sp. Nov., a novel non-saccharolytic anaerobe from the rumen that ferments pyruvate and amino acids, forms caproate and utilizes acetate and propionate. International Journal of Systematic and Evolutionary Microbiology 53:965–970.

33. Wallace RJ, Chaudhary LC, Miyagawa E, McKain N, Walker ND. 2004. Metabolic properties of E*eubacterium pyruvativorans*, a ruminal ‘hyper-ammonia-producing’ anaerobe with metabolic properties analogous to those of *Clostridium kluyveri*. Microbiology 150:2921–2930.

34. Jeon BS, Choi O, Um Y, Sang B-I. 2016. Production of medium-chain carboxylic acids by *Megasphaera* sp. Mh with supplemental electron acceptors. Biotechnology for Biofuels 9:129.

35. Holdeman LV, Cato EP, Moore WEC. 1967. Amended description of *Ramibacterium alactolyticum* prévot and taffanel with proposal of a neotype strain1. International Journal of Systematic and Evolutionary Microbiology 17:323–341.

36. Werner JJ, Garcia ML, Perkins SD, Yarasheski KE, Smith SR, Muegge BD, Stadermann FJ, DeRito CM, Floss C, Madsen EL, Gordon JI, Angenent LT. 2014. Microbial community dynamics and stability during an ammonia-induced shift to syntrophic acetate oxidation. Applied and Environmental Microbiology 80:3375–3383.

37. Allaart MT, Fox BB, Nettersheim IHMS, Pabst M, Sousa DZ, Kleerebezem R. 2023. Physiological and stoichiometric characterization of ethanol-based chain elongation in the absence of short-chain carboxylic acids. Science Reports 13:17370.

38. Katano Y, Fujinami S, Kawakoshi A, Nakazawa H, Oji S, Iino T, Oguchi A, Ankai A, Fukui S, Terui Y, Kamata S, Harada T, Tanikawa S, Suzuki K-i, Fujita N. 2012. Complete genome sequence of *Oscillibacter valericigenes* Sjm18-20(t) (=NBRC 101213(t)). Standards in Genomic Sciences 6:406–414.

39. Esquivel-Elizondo S, Bağcı C, Temovska M, Jeon BS, Bessarab I, Williams RBH, Huson DH, Angenent LT. 2021. The isolate Caproiciproducens sp. 7d4c2 produces *n*-caproate at mildly acidic conditions from hexoses: Genome and RBOX comparison with related strains and chain-elongating bacteria. Frontiers in Microbiology 11.

40. Spormann AM. 2024. Principles of microbial metabolism and metabolic ecology. Springer, Cham.

41. Kerfeld CA, Aussignargues C, Zarzycki J, Cai F, Sutter M. 2018. Bacterial microcompartments. Nature Reviews Microbiology 16:277–290.

42. Held M, Quin MB, Schmidt-Dannert C. 2013. EUT bacterial microcompartments: Insights into their function, structure, and bioengineering applications. Journal of Molecular Microbiology and Biotechnology 23:308–320.

43. Heldt D, Frank S, Seyedarabi A, Ladikis D, Parsons Joshua B, Warren Martin J, Pickersgill Richard W. 2009. Structure of a trimeric bacterial microcompartment shell protein, ETUB, associated with ethanol utilization in C*clostridium kluyveri*. Biochemical Journal 423:199–207.

44. Seedorf H, Fricke WF, Veith B, Brüggemann H, Liesegang H, Strittmatter A, Miethke M, Buckel W, Hinderberger J, Li F, Hagemeier C, Thauer RK, Gottschalk G. 2008. The genome of C*clostridium kluyveri*, a strict anaerobe with unique metabolic features. Proceedings of the National Academy of Sciences 105:2128–2133.

45. Contreras-Dávila CA, Carrión VJ, Vonk VR, Buisman CNJ, Strik DPBTB. 2020. Consecutive lactate formation and chain elongation to reduce exogenous chemicals input in repeated-batch food waste fermentation. Water Research 169:115215.

46. Srinivasan S, Kim HS, Kim MK, Lee M. 2012. *Pseudoclavibacter caeni* sp. Nov., isolated from sludge of a sewage disposal plant. International Journal of Systematic and Evolutionary Microbiology 62:786–790.

47. Coma M, Vilchez-Vargas R, Roume H, Jauregui R, Pieper DH, Rabaey K. 2016. Product diversity linked to substrate usage in chain elongation by mixed-culture fermentation. Environmental Science & Technology 50:6467–6476.

48. Vasudevan D, Richter H, Angenent LT. 2014. Upgrading dilute ethanol from syngas fermentation to *n*-caproate with reactor microbiomes. Bioresource Technology 151:378–382.

49. Usack JG, Angenent LT. 2015. Comparing the inhibitory thresholds of dairy manure co-digesters after prolonged acclimation periods: Part 1 – performance and operating limits. Water Res 87:446–457.

50. RStudio Team. 2016. Rstudio: Integrated development environment for r. RStudio, Inc., Boston, MA.

51. Regueiro L, Spirito CM, Usack JG, Hospodsky D, Werner JJ, Angenent LT. 2015. Comparing the inhibitory thresholds of dairy manure co-digesters after prolonged acclimation periods: Part 2 – correlations between microbiomes and environment. Water Research 87:458–466.

52. Bolyen E, Rideout JR, Dillon MR, Bokulich NA, Abnet CC, Al-Ghalith GA, Alexander H, Alm EJ, Arumugam M, Asnicar F, Bai Y, Bisanz JE, Bittinger K, Brejnrod A, Brislawn CJ, Brown CT, Callahan BJ, Caraballo-Rodríguez AM, Chase J, Cope EK, Da Silva R, Diener C, Dorrestein PC, Douglas GM, Durall DM, Duvallet C, Edwardson CF, Ernst M, Estaki M, Fouquier J, Gauglitz JM, Gibbons SM, Gibson DL, Gonzalez A, Gorlick K, Guo J, Hillmann B, Holmes S, Holste H, Huttenhower C, Huttley GA, Janssen S, Jarmusch AK, Jiang L, Kaehler BD, Kang KB, Keefe CR, Keim P, Kelley ST, Knights D, Koester I, Kosciolek T, Kreps J, Langille MGI, Lee J, Ley R, Liu Y-X, Loftfield E, Lozupone C, Maher M, Marotz C, Martin BD, McDonald D, McIver LJ, Melnik AV, Metcalf JL, Morgan SC, Morton JT, Naimey AT, Navas-Molina JA, Nothias LF, Orchanian SB, Pearson T, Peoples SL, Petras D, Preuss ML, Pruesse E, Rasmussen LB, Rivers A, Robeson MS, Rosenthal P, Segata N, Shaffer M, Shiffer A, Sinha R, Song SJ, Spear JR, Swafford AD, Thompson LR, Torres PJ, Trinh P, Tripathi A, Turnbaugh PJ, Ul-Hasan S, van der Hooft JJJ, Vargas F, Vázquez-Baeza Y, Vogtmann E, von Hippel M, Walters W, Wan Y, Wang M, Warren J, Weber KC, Williamson CHD, Willis AD, Xu ZZ, Zaneveld JR, Zhang Y, Zhu Q, Knight R, Caporaso JG. 2019. Reproducible, interactive, scalable and extensible microbiome data science using QIIME 2. Nature Biotechnology 37:852-857.

53. Virtanen P, Gommers R, Oliphant TE, Haberland M, Reddy T, Cournapeau D, Burovski E, Peterson P, Weckesser W, Bright J, van der Walt SJ, Brett M, Wilson J, Millman KJ, Mayorov N, Nelson ARJ, Jones E, Kern R, Larson E, Carey CJ, Polat İ, Feng Y, Moore EW, VanderPlas J, Laxalde D, Perktold J, Cimrman R, Henriksen I, Quintero EA, Harris CR, Archibald AM, Ribeiro AH, Pedregosa F, van Mulbregt P, Vijaykumar A, Bardelli AP, Rothberg A, Hilboll A, Kloeckner A, Scopatz A, Lee A, Rokem A, Woods CN, Fulton C, Masson C, Häggström C, Fitzgerald C, Nicholson DA, Hagen DR, Pasechnik DV, Olivetti E, Martin E, Wieser E, Silva F, Lenders F, Wilhelm F, Young G, Price GA, Ingold G-L, Allen GE, Lee GR, Audren H, Probst I, Dietrich JP, Silterra J, Webber JT, Slavič J, Nothman J, Buchner J, Kulick J, Schönberger JL, de Miranda Cardoso JV, Reimer J, Harrington J, Rodríguez JLC, Nunez-Iglesias J, Kuczynski J, Tritz K, Thoma M, Newville M, Kümmerer M, Bolingbroke M, Tartre M, Pak M, Smith NJ, Nowaczyk N, Shebanov N, Pavlyk O, Brodtkorb PA, Lee P, McGibbon RT, Feldbauer R, Lewis S, Tygier S, Sievert S, Vigna S, Peterson S, More S, Pudlik T, Oshima T, Pingel TJ, Robitaille TP, Spura T, Jones TR, Cera T, Leslie T, Zito T, Krauss T, Upadhyay U, Halchenko YO, Vázquez-Baeza Y, SciPy C. 2020. Scipy 1.0: Fundamental algorithms for scientific computing in python. Nature Methods 17:261-272.

54. Andrews S. 2010. Fastqc: A quality control tool for high throughput sequence data. Babraham Bioinformatics, Babraham Institute, Cambridge, United Kingdom.

55. Bolger AM, Lohse M, Usadel B. 2014. Trimmomatic: A flexible trimmer for illumina sequence data. Bioinformatics 30:2114–2120.

56. Buchfink B, Xie C, Huson DH. 2015. Fast and sensitive protein alignment using diamond. Nature methods 12:59.

57. Huson DH, Beier S, Flade I, Górska A, El-Hadidi M, Mitra S, Ruscheweyh H-J, Tappu R. 2016. Megan community edition-interactive exploration and analysis of large-scale microbiome sequencing data. PLoS computational biology 12.

58. Li D, Liu C-M, Luo R, Sadakane K, Lam T-W. 2015. Megahit: An ultra-fast single-node solution for large and complex metagenomics assembly via succinct de Bruijn graph. Bioinformatics 31:1674–1676.

59. Parks DH, Imelfort M, Skennerton CT, Hugenholtz P, Tyson GW. 2015. Checkm: Assessing the quality of microbial genomes recovered from isolates, single cells, and metagenomes. Genome research 25:1043–1055.

60. Seemann T. 2014. Prokka: Rapid prokaryotic genome annotation. Bioinformatics 30:2068–2069.

61. Finn RD, Bateman A, Clements J, Coggill P, Eberhardt RY, Eddy SR, Heger A, Hetherington K, Holm L, Mistry J. 2014. Pfam: The protein families database. Nucleic acids research 42:D222–D230.

62. Eddy SR. 2011. Accelerated profile hmm searches. PLOS Computational Biology 7:e1002195.

63. Sutter M, Kerfeld CA. 2022. BMC caller: A webtool to identify and analyze bacterial microcompartment types in sequence data. Biology Direct 17:9.

