## Supporting Information for "The gases H_2_ and O_2_ in open-culture reactors influence the performance and microbiota of chain elongation into *n*-caproate and *n*-caprylate"

Running Title: The microbiome of C8-producing reactors

Authors: Catherine M. Spirito[1,2*], Timo N. Lucas[3*], Sascha Patz[3], Byoung Seung Jeon[4], Jeffrey J. Werner[5], Lauren H. Trondsen[1], Juan J. Guzman[1], Daniel H. Huson[3], Largus T. Angenent[1,4,6,7,8]**

[1] Department of Biological and Environmental Engineering, Cornell University, Riley-Robb Hall, Ithaca, NY 14853 Ithaca, NY 14853, USA

[2] Office of Undergraduate Research, University of Maryland, College Park, MD, 20742, USA

[3] Institute for Bioinformatics and Medical Informatics, University of Tübingen, Sand 14, 72076 Tübingen, Germany

[4] Department of Geosciences, University of Tübingen, Schnarrenbergstr. 94-96, 72076 Tübingen, Germany

[5] Chemistry Department, SUNY-Cortland, Bowers Hall, Cortland, NY 13045, USA

[6] AG Angenent, Max Planck Institute for Biology Tübingen, Max Planck Ring 5, 72076 Tübingen, Germany

[7] Department of Biological and Chemical Engineering, Aarhus University, Gustav Wieds Vej 10D, 8000 Aarhus C, Denmark

[8] The Novo Nordisk Foundation CO2 Research Center (CORC), Aarhus University, Gustav Wieds Vej 10C, 8000 Aarhus C, Denmark

*Indicates shared first author status

**Correspondence to:

**Largus T. Angenent**


Number of pages: 14; Number of figures: 3; Number of tables: 3

**Supporting Info - Methods**

*Continuously Fed Reactor System*

At the start of the study, the reactors were inoculated with 10% by volume (~500 mL) of yeast fermentation beer reactor broth from a reactor fed with ethanol-rich yeast fermentation beer and operated as an anaerobic sequencing batch reactor for a period of approximately five years *prior* to the time the inoculum was collected (1, 2).

*Gas Sparging Setup*

Gas exit lines from the top of the reactor led to a condensation trap, bubbler, and then a gas flow meter (Calibrated Instruments Inc., Ritter MilliGas Counter Series MGC-1 V3.1, Hawthorne, NY). Stainless-steel lines were setup to sparge the reactor broth with either $N_2$, $H_2$, or both gases combined. The lines ran from tanks of $N_2$ or $H_2$ gas (pressure of tank set at 40 psi) to the reactor and entered the reactor near the reactor base. One-way check valves located near the reactor base were used to ensure uni-directional flow into the reactor. Gas flow meters (Cole-Parmer 65-mm Correlated Flowmeter, Part No. EX-03216-02), as well as needle valves, were used to control the flow rate of the gases into the reactor system. The gas flow rate entering the reactors was not measured. The gas flow rates leaving the reactors were measured and are reported in **Table 1** (in the main text).

*Pertraction System*

For product extraction, we used a setup similar to the one previously described by Agler et al. (1). Forward and backward membrane contactors (8.1 $m^2$ each, Membrana Liqui-Cel 4x13, X50 Membrane, Charlotte, NC, USA) were connected to the reactor setup (**Fig. S1**). A peristaltic pump (Cole Parmer 7553-30) was used to recirculate reactor broth at a flow rate of 48 mL $min^{-1}$ through the shell-side of the forward contactor. Broth was removed from near the top of the reactor, passed through a filter (McMaster Carr High-Pressure Stainless Steel Y-Strainer, 1/2 NPT Female, 100 Mesh) to remove particulate matter, and then passed through the contactor and back to the reactor. The filter was periodically cleaned to remove the built-up particulate matter, on an approximately monthly basis. A mineral oil solvent with 30 g $L^{-1}$ tri-*n*-octylphosphine oxide (TOPO) (Sigma Aldrich, St. Louis, MO, USA) was continuously circulated at a flow rate of ~30 mL $min^{-1}$ (Cole Parmer 7553-30) through the lumen side of both the forward and backward membrane contactors. The purpose of the solvent was to primarily extract the more hydrophobic medium-chain carboxylates (as compared to the short-chain carboxylates) from the reactor broth. A well-mixed alkaline extraction solution was recycled at a flow rate of ~30 mL $min^{-1}$ (Cole Parmer 7553-30) through the shell-side of the backward contactor. 0.3M sodium borate was initially used to buffer the extraction solution. The pH of the alkaline extraction solution was maintained at ~pH 9 *via* automated additions of 5 M NaOH using a pH controller and a corresponding base-addition pump.

**Supporting Information – CSV Files**

The following csv files are provided with the supporting info:

- RBOX protein counts for reactor 1, timepoint 1
    - File name: r1t1_rbox_counts.csv
- RBOX protein counts for reactor 2, timepoint 1
    - File name: r2t1_rbox_counts.csv
- Bacterial microcompartments (BMCs) identified in reactor metagenomics samples using BMC caller.
    - File name: BMC_from_BMC_caller_tool.csv

**Supporting Info - Tables**

**Table S1.** Percent hydrogen (by volume) measured in the headspace of the three reactors. Mean and s.d. values are reported per main period.

| | Period | $H_2$ (%) |
|---|---|---|
| Reactor 1 | 1 | 9.9 ± 5.2 |
| | 2 | 1.8 ± 0.6 |
| | 3 | 0.7 ± 0.2 |
| Reactor 2 | 1 | 1.8 ± 1.9 |
| | 2 | 18 ± 19 |
| | 3 | 13 ± 5.3 |
| Reactor 3 | 1 | 31 ± 9.6 |
| | 2 | 20 ± 14 |
| | 3 | 7.3 ± 4.6 |

**Table S2.** Effluent ethanol and carboxylate concentrations per period. Mean and standard error reported. Periods 1, 2, and 3 are abbreviated as P1, P2, and P3 in this Table.

| | | Ethanol (mM) | Acetate (mM) | *n*-Butyrate (mM) | *n*-Caproate (mM) | *n*-Caprylate (mM) |
|---|---|---|---|---|---|---|
| Reactor 1 | P1 | 47.4 ± 3.9 | 3.97 ± 0.44 | 2.22 ± 0.11 | 2.26 ± 0.12 | 1.68 ± 0.12 |
| | P2 | 61.0 ± 4.01 | 4.12 ± 0.39 | 5.26 ± 0.28 | 3.77 ± 0.17 | 0.98 ± 0.08 |
| | P3 | 73.9 ± 4.12 | 7.92 ± 1.11 | 5.88 ± 0.54 | 3.75 ± 0.21 | 0.77 ± 0.07 |
| Reactor 2 | P1 | 29.1 ± 4.25 | 9.15 ± 0.90 | 4.20 ± 0.27 | 3.52 ± 0.22 | 0.97 ± 0.13 |
| | P2 | 69.8 ± 7.54 | 11.7 ± 1.71 | 4.67 ± 0.34 | 3.68 ± 0.47 | 1.84 ± 0.34 |
| | P3 | 115 ± 9.09 | 13.6 ± 1.32 | 2.33 ± 0.15 | 2.06 ± 0.12 | 0.54 ± 0.04 |
| Reactor 3 | P1 | 166 ± 9.69 | 2.55 ± 0.40 | 4.07 ± 0.53 | 1.58 ± 0.19 | 0.38 ± 0.04 |
| | P2 | 119 ± 11.2 | 1.08 ± 0.18 | 2.39 ± 0.27 | 1.74 ± 0.17 | 0.48 ± 0.05 |
| | P3 | 98.1 ± 5.86 | 3.40 ± 0.54 | 4.59 ± 0.47 | 4.05 ± 0.26 | 1.04 ± 0.07 |

**Table S3:** Components of stainless-steel reactor system. Vendor, catalog number, and a brief description provided.

| Vendor | Catalog Number | Description | Purpose | Detailed purpose |
|---|---|---|---|---|
| McMaster Carr | 52245K535 | 316 stainless steel compression tube fitting, adapter, for 3/8" tubing, 1/4" pipe | Connections | Effluent port |
| McMaster Carr | 52245K824 | Stainless steel compression tube fitting adapter for 3/8 NPT, 3/8" tube OD | Connections | System recycle |
| McMaster Carr | 52245K825 | Stainless steel compression tube fitting adapter for 1/2 NPT, 3/8" tube OD | Connections | Internal recycle |
| Swagelok | MS-PTS-6 | Swak Pipe Thread Sealant, 6 cc | Connections | Sealing |
| MMC | 7098K25 | Slim Spade-Terminal Relay SPDT, 12 DC Control Voltage | Electrical | Liquid level |
| McMaster Carr | 7527K53 | Terminal block | Electrical | Electrical connections (on plywood) |
| McMaster Carr | 7098K23 | Socket with spring clamps (for relay) | Electrical | Liquid level |
| McMaster Carr | 2779K5 | LED Panel mounting indicating light | Electrical | liquid level |
| ebay | | Power strips | Electrical | Electrical connections (on plywood) |
| ebay | | 12 V DC power supply | Electrical | Liquid level |
| MMC | 7527K59 | Jumpers (pack of 25) | Electrical | Liquid level |
| MMC | 7527K836 | Cover for terminal block | Electrical | Electrical connections (on plywood) |
| MMC | 70355K85 | Power Cord with Three-Blade Plug 18 Gauge Wire, 6' Long SPT-1 Cord | Electrical | Control board |
| MMC | 7196K41 | Straight-Blade Three-Blade Female Connector, NEMA 5-15 | Electrical | Control board |
| MMC | 8054T15 | Stranded Wire 300V AC, 18 Gauge, 200 ft, Black | Electrical | Control board |
| MMC | 8054T15 | Stranded Wire 300V AC, 18 Gauge, 200 ft, Red | Electrical | Control board |
| MMC | 8054T15 | Stranded Wire 300V AC, 18 Gauge, 200 ft, White | Electrical | Control board |
| Quantumflow Tech | | 4x13 liqui-cel membrane contactors | Extraction | Extraction |

| | | | | |
|---|---|---|---|---|
| McMaster Carr | 51525K236 | Plastic Quick-Turn (Luer Lock) Coupling, Polypropylene, Female x Male Thread, 1/4"-28 UNF Thread (Pack of 10) | Filter | Filter |
| McMaster Carr | 44205K21 | Disposable Water Filter (5 microns) | Filter | Filter |
| Lowes | 253209 | Toilet Flange Extender and Complete Spacer (2 spacers) | Filter | Strainer |
| Lowes | 23493 | 4-in PVC Cap Fitting | Filter | Strainer |
| Lowes | 126605 | 4 1/2 in dia Stainless steel basket | Filter | Strainer |
| Lowes | 137639 | 1/4" Bolts | Filter | Strainer |
| Lowes | 67340 | 1/4" Nuts | Filter | Strainer |
| Lowes | 61814 | 1/4" Washers | Filter | Strainer |
| MMC | 5670K85 | Type 303 Stainless Steel Barbed Tube Fitting, 1/4 NPT, 3/8" ID tube | Filter | Strainer |
| MMC | 4452K672 | Locknut 1/4 NPT | Filter | Strainer |
| MMC | 91525A145 | Large diameter flat washer for 1/2" screw size, pack of 10 | Filter | Strainer |
| Airgas | HY HP300 | $H_2$ gas tank | Gas | |
| Ritter US LLC | | Series MGC-1 Milli Gas Counter V 3.1 | Gas | Gas measurement |
| McMaster Carr | 89785K43 | 316 stainless steel tubing 3/8" OD | Lines | Liquid and gas lines |
| McMaster Carr | 89785K122 | 316 stainless steel tubing, 1/4" OD | Lines | pH lines? |
| McMaster Carr | 4066K41 | Stainless steel pressure gauge (0-30 psi) | Lines | Pressure measurements |
| Omega | LV-11 | Liquid level switch, stainless steel | Liquid level | Liquid level |
| Omega | PHE-7353-15 | pH probe | pH | Probe |
| Cole Parmer | YO-56705-00 | pH controller | pH | Controller |
| Cole Parmer | 07554-80 | Pump drive | Pumps | |
| Cole Parmer | 77250-62 | High pressure pump head | Pumps | |
| Cole Parmer | S-95564-24 | High pressure tubing | Pumps | |
| Digikey | AE9869-ND | CABLE DB9M-DB9M 2M (for pumps) | Pumps | For back connection |

| | | | | |
|---|---|---|---|---|
| McMaster Carr | 44635K644 | 316 stainless steel unthreaded pipe size 4, 6' length | Reactor | Reactor body |
| McMaster Carr | 44695K39 | 316 stainless steel low pressure unthreaded flange | Reactor | Reactor body |
| McMaster Carr | 44695K119 | 316 stainless steel cap | Reactor | Not needed |
| McMaster Carr | 45555K121 | Reducing coupling; pipe size 4 to 1 1/2" | Reactor | Bottom |
| McMaster Carr | 9472K49 | All-purpose gasket | Reactor | |
| McMaster Carr | 91236A804 | Znc-Pltd STL Low-Strength Hex Head Cap Screw 5/8"-11 Thread, 2-1/2" Length, packs of 10 | Reactor | Reactor flange |
| McMaster Carr | 4452K112 | Stainless steel coupling, 1/4 NPT | Reactor | Gas outlet, effluent, extra port |
| McMaster Carr | 4452K138 | Stainless steel half coupling, 1/8 NPT | Reactor | Temp |
| McMaster Carr | 4452K114 | Stainless steel coupling for 1/2 pipe size | Reactor | |
| McMaster Carr | 52245K539 | Stainless steel compression tube fitting adapter for 1/2 NPT, 1/2" tube OD | Reactor | Not needed |
| McMaster Carr | 90108A035 | Washers for 5/8" screw size, pack of 25 | Reactor | Reactor flange |
| McMaster Carr | 95462A533 | Nuts for 5/8" screw size, pack of 50 | Reactor | Reactor flange |
| McMaster Carr | 4942K7 | Plastic sight glass (window diameter 2 13/16"), for 3" tube OD | Reactor | Window |
| McMaster Carr | 50485K165 | Adapter, quick clamp to tube | Reactor | Window |
| McMaster Carr | 4322K155 | Clamp | Reactor | Window |
| McMaster Carr | 43315K27 | PTFE gasket for reactor window | Reactor | Window |
| McMaster Carr | 4452K213 | Half coupling 3/4" pipe size | Reactor | pH port |

| | | | | |
|---|---|---|---|---|
| McMaster Carr | 4452K112 | Coupling for 1/4 pipe size | Reactor | Effluent, extra port |
| McMaster Carr | 4452K139 | Half coupling 1/4" pipe size | Reactor | Gas line |
| McMaster Carr | 4452K111 | Coupling for 1/8 pipe size | Reactor | Temp |
| McMaster Carr | 4942K3 | Plastic sight (reactor window) for tube OD 2" | Reactor | Window |
| McMaster Carr | 43315K25 | PTFE gasket for reactor window (2") | Reactor | Window |
| McMaster Carr | 50485K163 | quick clamp tube fittings for tube OD 2" | Reactor | Window |
| McMaster Carr | 4322K153 | Clamp (2") | Reactor | Window |
| McMaster Carr | 47865K41 | Brass ball valve 1/4 NPT, female x male | Reactor | Not needed? |
| McMaster Carr | 7768K22 | Brass check valve (female inlet x male outlet, 1/4 pipe size) | Reactor | |
| McMaster-Carr | 4322K714 | Clamp for reactor window (bolted) | Reactor | |
| McMaster Carr | 44635K832 | 316 stainless steel unthreaded pipe size 4, 3' length | Reactor | Body |
| McMaster Carr | 45555K122 | Reducing coupling; pipe size 4 to 2" | Reactor | |
| McMaster Carr | 50485K521 | Stainless Steel Quick-Clamp Tube Fitting, Female Pipe Adapter for 1" Tube OD, 1" NPT Pipe | Reactor | |
| McMaster Carr | 4322K152 | Stainless Steel Quick-Clamp Tube Fitting, Wing-Nut Clamp for 1 &1-1/2" Tube OD, 1.984" Flange | Reactor | |
| McMaster Carr | 4509K13 | Buna-N Gasket for Sanitary Tube Fitting for 1" Tube OD | Reactor | |
| McMaster | 4452K125 | Stainless steel square head plug for 3/4 pipe size | Reactor | Temporary |
| McMaster Carr | 4452K113 | Stainless steel coupling, 3/8 NPT | Reactor | System recycle out |
| McMaster Carr | 4452K138 | Half stainless steel coupling, 1/8 NPT | Reactor | For liquid level |
| McMaster Carr | 4452K421 | 45 deg. stainless steel elbow, female, 1/8 NPT | Reactor | For liquid level |

| | | | | |
|---|---|---|---|---|
| McMaster Carr | 4548K111 | Stainless steel thread nipple, 1/8 NPT (close) | Reactor | For liquid level |
| MMC | 4452K111 | Stainless steel 1/8 NPT coupling | Reactor | For liquid level |
| MMC | 4816K153 | Stainless steel pipe (1/8 NPT, threaded both ends, 20" length) | Reactor | For liquid level |
| MMC | 4816K121 | Stainless steel pipe (1/8 NPT, threaded both ends, 18" length) | Reactor | For liquid level |
| MMC | 4548K115 | Stainless steel pipe (1/8 NPT, threaded both ends, 3" length) | Reactor | For liquid level |
| Home Depot | ERZ782478W-4 | Commercial shelving unit; 77 in W x 78 in H x 24 in D for reactor stand | Stand | |
| Lowes | | Paint | Stand | |
| Lowes | | Primer | Stand | |
| Home Depot | | Plywood | Stand | |
| McMaster-Carr | 98750A214 | 5/8" threaded rod for reactor stand/setup | Stand | |
| McMaster Carr | 90108A035 | Washers for 5/8" screw size, pack of 25 | Stand | |
| McMaster Carr | 95462A533 | Nuts for 5/8" screw size, pack of 50 | Stand | |
| McMaster Carr | 90322A335 | 5/8" thread steel rod for reactor stand (6 ft) | Stand | |
| MMC | 35765K277 | Heat sheet (3"x12") | Temp | Heater |
| Omega | HSTC-TT-K-20S-120 | Type K Hermetically sealed thermocouple, 3 m (120"), stripped leads | Temp | For under heater |
| MMC | 5556K39 | Rigid high temperature fiberglass pipe insulation | Temp | Insulation |
| MMC | 45325K114 | Plastic jacketing for pipe insulation | Temp | Insulation |
| MMC | 29695T123 | Cinching strap | Temp | Insulation |
| ebay | | Heat transfer paste, 1 oz | Temp | Heater |
| Amazon | | Dual Digital Display PID Temperature Controller SSR(2 Alarms) | Temp | Temp control |
| Omega | TJ36-CASS-18U-6 | Thermocouple, type K, stainless steel, 6 " length, 1/8" diameter | Temp | For reactor |
| Omega | SSLK-18-18 | 1/8 x 1/8 compression fitting | Temp | For reactor temp |

| | | | | |
|---|---|---|---|---|
| MMC | 3869K35 | Thermocouple and RTD Connector Male, 400 Deg F, Flat-Pin, Type K, Yellow | Temp | Heater |
| MMC | 3869K34 | Thermocouple and RTD Connector Female, 400 Deg F, Flat-Pin, Type K, Yellow | Temp | Heater |

**Supporting Info - Figures**

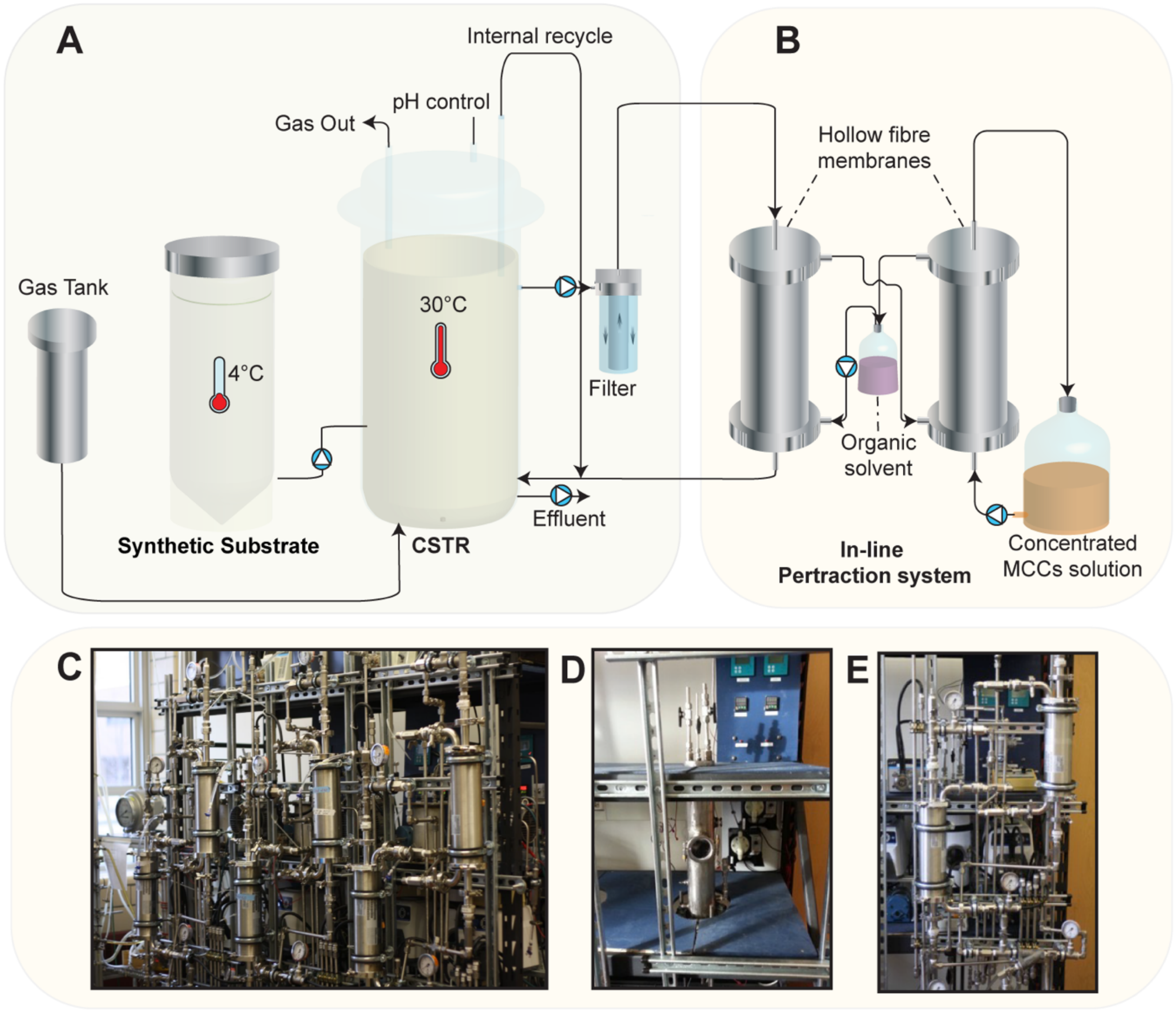

**Figure S1.** Diagram showing setup of each reactor. Gas was sparged into the bottom of the reactor during Periods 2 and 3 (**A**). Pertraction system setup with membrane contactors, oil, and alkaline extraction solution (**B**). Photos of the setup of the in-house constructed, three stainless-steel reactors: the membrane contactors and stainless-steel lines of the pertraction system are shown (**C**); stainless steel CSTR and electrical control board (**D**) and single reactor with pertraction system (**E**).

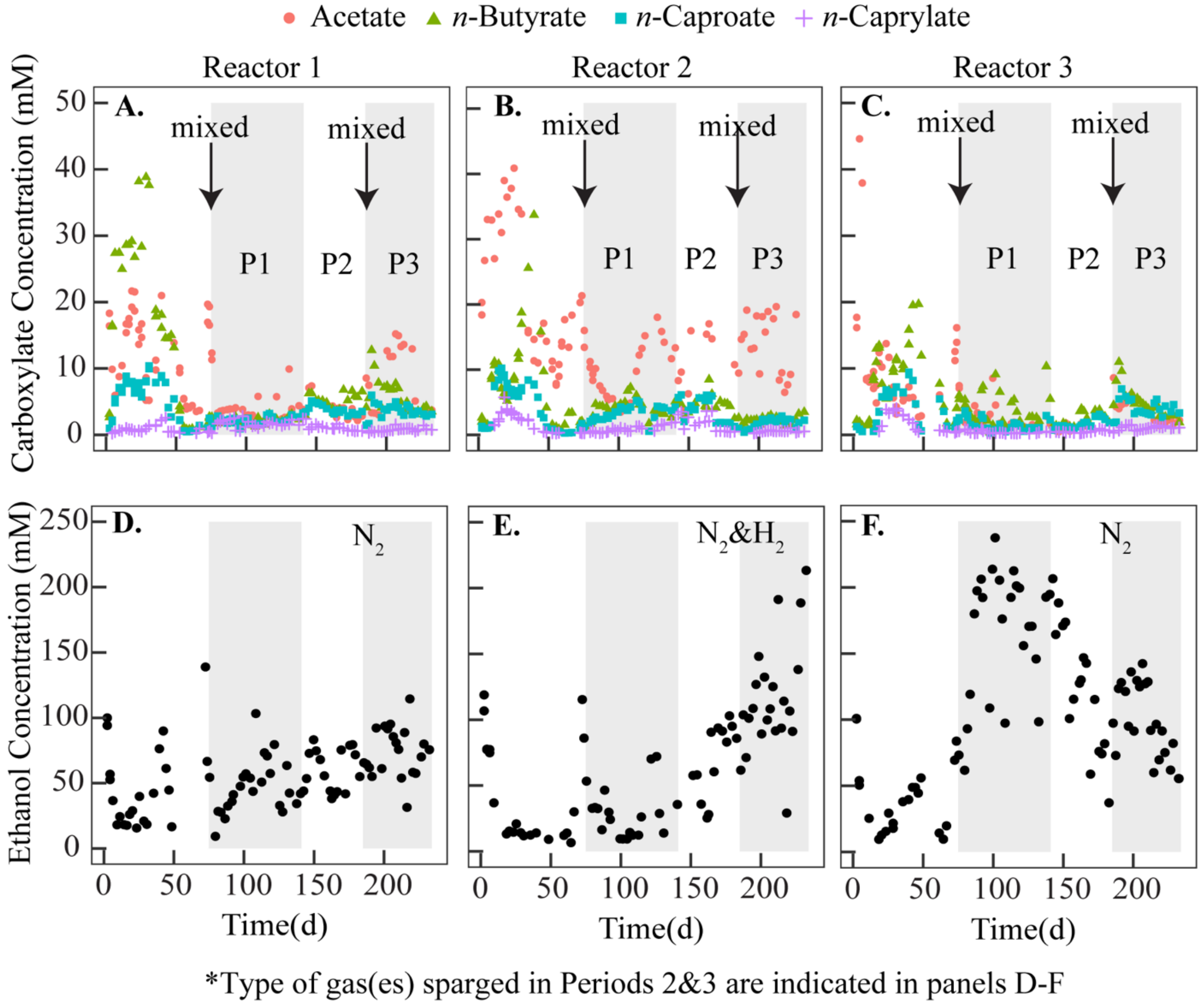

**Figure S2.** Effluent carboxylate concentrations throughout the operating period for Reactor 1 (**A**), Reactor 2 (**B**), and Reactor 3 (**C**). The two-time points at the start of Periods 1 and 3, when the biomass from all the reactors was mixed, are indicated on these panels. Effluent ethanol concentrations throughout the operating period for Reactor 1 (**D**), Reactor 2 (**E**), and Reactor 3 (**F**) are also shown. The type of gases that were sparged into these reactors during Periods 2 &3 are indicated in these panels. Dark shading in panels **A-F** indicate Periods 1, 2, and 3 (P1, P2, and P3), which are the main periods reported on in this text.

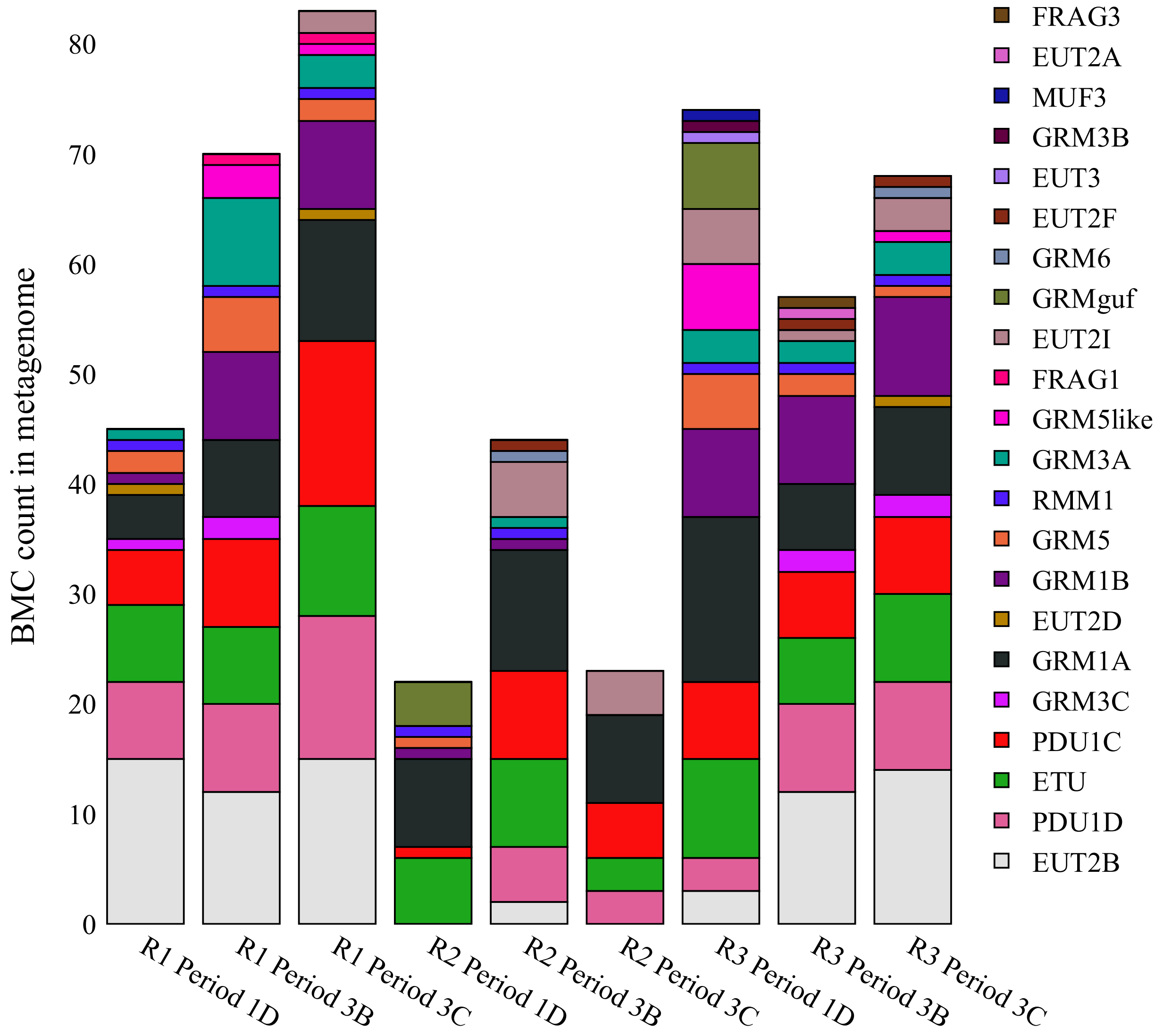

**Figure S3**. Count of bacterial microcompartment (BMC) shell proteins in annotated metagenomes. Each sampling point is represented by a distinct bar and the bar heights are proportional to the number of microcompartments found. The bar segments are colored by the microcompartment's type. R1-3 are Reactors 1-3. The periods are explain in the methods of the main text.

### References

1. **Agler MT, Spirito CM, Usack JG, Werner JJ, Angenent LT.** 2012. Chain elongation with reactor microbiomes: Upgrading dilute ethanol to medium-chain carboxylates. Energy and Environ Science **5:**8189-8192.
2. **Ge S, Usack JG, Spirito CM, Angenent LT.** 2015. Long-term *n*-caproic acid production from yeast-fermentation beer in an anaerobic bioreactor with continuous product extraction. Environtal Science and Technology **49:**8012-8021.
